# Analysis of the Neurotropic and Anti-inflammatory Effects of Neuroelectrode Functionalisation with a Heparan Sulphate Mimetic

**DOI:** 10.1101/2021.05.11.443655

**Authors:** Catalina Vallejo-Giraldo, Idir Ouidja Mohand, Minh Bao Huynh, Alexandre Trotier, Katarzyna Krukiewicz, Anuradha R Pallipurath, Andrea Flannery, Michelle Kilcoyne, Abhay Pandit, Dulce Papy-Garcia, Manus Jonathan Paul Biggs

## Abstract

Further in the search for biomimicry of the properties analogous to neural tissues, and with an ultimate goal of mitigating electrode deterioration via reactive host cell response and glial scar formation, the bio-functionalisation of PEDOT:PTS neural coating is here presented using a heparan mimetic termed (HM) F6. A sulphated mimetic polyanion, with a potential role in neuromodulation in neurodegenerative diseases, and used here for the first time as neural coating.

This work acts as a first step towards the use of HM biological dopants, to enhance neuroelectrode functionality, to promote neural outgrowth and to maintain minimal glial scar formation in vitro at the neural-interface. Further, this study opens new possibilities for the evaluation of glycan mimetics in neuroelectrode functionalisation.

## 1. Introduction

Chronic biocompatibility of an implantable neural device can be enhanced through biomimicry strategies to recapitulate the physicomechanical and biochemical properties of neural tissues, with an ultimate goal of mitigating electrode deterioration via a reactive host response [1–5]. The single most detrimental factor in chronic electrode stability is mediated through an adverse peri-electrode tissue response characterised by glial scar formation and electrode encapsulation, causing a reduction in signal strength and local death of adjacent neurons [5–9]. In order to develop meaningful neuromodulation systems and stable brain-machine interfaces there is a clinical need to reduce tissue encapsulation and improve long-term neuroelectrode function through enhanced integration *in situ* [10].

Conducting polymers acting as interfacial neuroelectrode coatings, provide excellent opportunities for functional modification due to their versatile chemistry and simple synthesis approaches [11–13] allowing the preparation of electroactive biointerfaces with tuned electrochemical [14, 15], topographical [16–18] and biochemical [19–21] properties.

Of particular interest, poly(3,4-ethylenedioxythiophene) (PEDOT) derived biomaterials have received considerable interest in the bioengineering of neural implants and devices due to their relatively high conductivity and stability properties [10, 22–24], as well as their superior biocompatibility properties, which can be tailored through the immobilisation of biological elements [2, 25–28] and anti-inflammatory drugs into the polymer matrix [20, 29–31].

In particular, the therapeutic potential of glycosaminoglycans (GAGs) - naturally occurring heteropolysaccharide molecules associated with core protein or lipid molecules, as bio-functional approaches to neurospecific biomaterials has been recently explored [28, 32–36]. This focus on GAG functionalisation has stemmed from the elucidation that GAG species play key roles in the extracellular matrix (ECM) as a major reservoir of growth factors [37–39] and as biological regulators of cellular growth, differentiation and regenerative processes [40–43].

Of particular interest in neural regeneration is the sulphated proteoglycan, heparan sulphate (HS) that through binding growth factors, plays a critical role in regulating inflammatory responses [43–49] and has shown marked efficacy as a therapeutic functionalisation approach within a broad spectrum of tissue engineering and implantable device applications [40, 50–60].

Accordingly, new avenues of research, focused on the development of synthetic heparan mimetic (HM) chemistries have been initiated to develop stable bioactive glycans for neural engineering applications [35, 61–64]. Of specific interest for the development of neuroelectrode glycofunctionalisation approaches is the formulation of HMs with inherent glycanase resistance, which can circumvent the prolong inflammation profile and perturbed remodeling of the ECM associated with degraded HS [49, 52, 65–67].

In this work, a PEDOT:PTS conducting polymer was doped with a HM termed F6 [44, 68], a polyanion molecule, structurally modified from polymeric dextran and shown to mediate internalisation and propagation of specific proteopathic proteins [60]. The effects of PEDOT:PTS:F6 functionalisation on the physical, electrochemical and biological properties of platinum (Pt) microelectrodes were subsequently evaluated *in vitro*.

## 2. Materials and Methods

### 2.1.1 Physical Characterisation

#### 2.1.1.1 Surface Morphology

Scanning electron microscopy (SEM) was carried out using a Hitachi S-4700 Cold Field Emission Gun Scanning Electron Microscope (CFE-SEM). The SEM images were taken using an accelerating voltage of 15 kV and spot current of 10 µA.

For roughness analysis, Atomic Force Microscopy (AFM) was performed as detailed in [5]. All measurements were taken on a Vico Dimension 3100 AFM using TESPA Tips (NanoWorld) (Si <8 nm tip radius, 42 N/m spring constant, 320 kHz nominal resonance frequency), in tapping mode over an area of 10 µm^2^ with a 0.5 - 1 Hz scan rate.

#### 2.1.1.2 Thickness Measurements

The thickness of the polymeric PEDOT:PTS and PEDOT:PTS:F6 coatings was measured using a Zygo Newview 100 surface profilometer controlled by MicroPlus software as detailed in [5]. Briefly, a pattern of bright and dark lines - fringes was created as incoming light was split from the limited region between the sample coating and the metal part of the electrode. This pattern difference was translated to calculate the height information.

### 2.1.2 Chemical Characterisation

#### 2.1.2.1 Contact Angle Measurement

Contact angles analysis with water droplets was performed using a custom made goniometer connected to an Infinity Camera 2 and operated with Infinity Analyse software. Contact angles were measured at three different points per sample in at least three samples per group. The images taken were calibrated using the measured width of the needle, and through the Infinity Analyse software a curve which was fitted to the arc of the drop to find the radius (R), as well as the length (L) of the base of the drop (from corner to corner).

The equation used to calculate the contact angle is shown below (***Equation 1***), where a (−) negative sign was used for hydrophilic surfaces and (+) sign was used for hydrophobic surfaces.

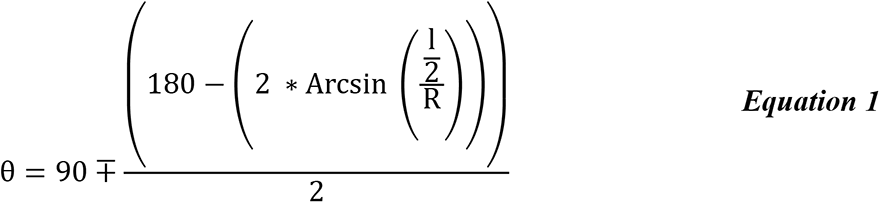

#### 2.1.2.2 Attenuated Total Reflectance Infra-Red Spectroscopy (ATR-IR)

Fourier transform mid infrared spectra in the range of 650-3600 cm^−1^, with a resolution of 4 cm^−1^ and four integrated scans, were collected on a diamond/ZnSe window using a Perkin Elmer Spectrum 400 fitted with an ATR reflectance attachment.

#### 2.1.2.3 Raman spectroscopy

Raman spectra were collected using the Renishaw Invia micro-Raman spectrometer (static acquisition centered at 800cm^−1^, 1 cm^−1^ resolution, 10s exposure time, 1% laser power, 785 nm laser, 4-5 µm spatial resolution, 12 µm penetration depth). Raman mapping was carried out using the same setup, where 81 spectra were collected at 5 µm intervals in a 60 × 60 µm grid. All spectra were normalised to account for variation in sample height on the intensity. Component DLS analysis was carried out using the Renishaw Wire 4.2 software to identify spectral contributions from different components.

#### 2.1.2.4 Ray Photoelectron Spectroscopy

X-ray photoelectron spectroscopy (XPS) spectra were acquired on a Kratos AXIS 165 spectrometer XPS system with X-Ray Gun mono Al Kα 1486.58 eV; 150 W (10 mA, 15kV), for all scans with the following parameters: sample temperature in a range of 20-30 oC with a pass energy of 160 eV for survey spectra and 20 eV for narrow regions and steps of 1 eV for survey and 0.05 eV for regions with dwell times of 50 ms and 100 ms for regions and sweeps for survey of ∼35, and for narrow regions of 6-40. The C1s line at 284.8 eV was used as charge reference. Spectra were collected in the normal way to the surface direction with an analysis area of 60 microns. XPS detection limit is estimated to be ∼0.1 at%. For the data processing, the construction and peak fitting of synthetic peaks in narrow region spectra was done using a Shirely type background and the synthetic peaks were of a mixed Gaussian-Lorenzian type. Relative sensitivity factors used are from CasaXPS library containing Scofield cross-sections.

### 2.1.3 Electrochemical Characterisation

#### 2.1.3.1 Preparation of PEDOT:PTS and PEDOT:PTS:F6 samples

The electrodeposition of poly(3,4-ethylenedioxythiophene):p-toluene sulfonate (PEDOT:PTS) was conducted under ambient conditions according to methods described previously [23]. Briefly, a solution of 0.05M EDOT (Sigma Aldrich, Ireland) and 0.1 M PTS (Sigma Aldrich, Ireland, 70,000 g mol^−1^ MW) was prepared in a 50/50 vol.% mixture of acetonitrile and water. The same prepared electrolyte with increasing concentrations of heparan sulphate mimetic F6, i.e. 1 µg ml^−1^, 10 µg ml^−1^, 50 µg ml^−1^, 100 µg ml^−1^ and 1000 µg ml^−1^ was used for the electrodeposition of the functionalised PEDOT:PTS:F6 coatings. F6 has a MW of 9900 g mol^−1^ (Da). The electrolyte solution was placed in an in-house fabricated electrochemical cell, connected to a Princeton Applied Research Potentiostat/Galvanostat model 2273 controlled with Power Suite software. Parylene-C insulated Platinum/Iridium concentric microelectrodes (MicroProbes for Life Science) and a platinum foil (Goodfellow) were used as the working electrode (WE) and counter-electrode (CE) respectively. A saturated 3 M KCl Ag/AgCl reference electrode (RE) (Bioanalytical Systems) was employed. Galvanostatic electrodeposition was performed and the efficiency of coating, i.e. the amount of polymer deposited on the electrodes, was controlled by the total charge passing during the electrodeposition. When the deposition was finalised, the pristine PEDOT:PTS coated electrodes were soaked in deionized water (DI) for 24 hours to remove excess of electrolyte and subsequently dried for use. The PEDOT:PTS:F6 coated electrodes were soaked in Hank’s balanced salt solution (HBSS) for two hours. For cell studies, pristine PEDOT:PTS and PEDOT:PTS:F6 were electrodeposited on electrodes with areas of 1.6 cm^2^ to facilitate *in vitro* manipulations.

To determine the total amount of F6 incorporated into PEDOT:PTS polymer, the electrochemical deposition was carried out by means of CH Instruments 400c Electrochemical Workstation equipped with time-resolved electrochemical quartz crystal microbalance in a standard three-electrode setup, employing CHI125 gold crystal as a working electrode, Ag/AgCl reference electrode and a platinum foil counter electrode.

The change of mass of the electrode resulting from the electrodeposition of pristine PEDOT:PTS or PEDOT:PTS:F6 coatings, respectively, was calculated based on frequency change (Δf) as given by the Sauerbrey equation (***Equation 2***) [69]:

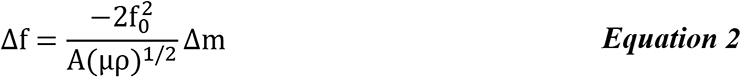

where f_0_ is the resonant frequency of the fundamental mode of the crystal, A is the area of the gold disk coated onto the crystal (0.205 cm^2^), ρ is the density of the crystal (2.648 g cm^−3^) and µ is the shear modulus of quartz (2.947·1011 g cm^−1^ s^−2^). For 8 MHz crystal, 1 Hz change in frequency corresponds to a mass change of 1.4 ng.

#### 2.1.3.2 Electrochemical Measurements

Cyclic voltammetry (CV) was performed as previously described in [5] using a Princeton Applied Research Potentiostat/Galvanostat model 2273 running with Power Suite software. Measurements were recorded in a custom-made electrochemical cell containing the Parylene-C insulated Platinum/Iridium concentric microelectrode as working electrode, an Ag/AgCl reference electrode (3 M KCl) (Bioanalytical Systems) and a platinum foil counter electrode (Goodfellow) in 0.01 M phosphate-buffered saline (PBS-1X). Cyclic voltammograms were run in the potential range from −0.8 V to 1 V at a scan rate of 0.1 V s^−1^. The charge storage capacity (CSC) was calculated by integrating the area enclosed by the voltammogram.

Electrical impedance spectroscopy (EIS) was performed using a Princeton Applied Research Potentiostat/Galvanostat model 2273 running with Power Suite software with a three-electrode set-up. The measurements were carried out at a frequency range of 0.1 Hz to 100 kHz with an AC sine wave of 40 mV amplitude applied with 0 V DC offset. The results were presented on Bode and Nyquist plots and compared to those of bare platinum/iridium microelectrodes. The data fitting analysis was performed using EIS Spectrum Analyser 1.0 software with the application of Powell algorithm.

The electrochemical stability of pristine PEDOT:PTS and PEDOT:PTS:F6 microelectrodes was done by performing cyclic voltammetry before and after 1000 stimulation cycles, corresponding to the cumulative stimulus used for the active release studies, biphasic potential pulse consisting of a 5-second application of a reduction potential (−0.5 V) followed by a 5-second application of an oxidative potential (+0.5 V). The percentage of loss in charge storage capacity was calculated by comparing the CSC of the 1st and 1000th cycles. All measurements were performed in triplicates; the results were expressed as a mean ± standard deviation.

### 2.1.4 Biological Characterisation

#### 2.1.4.1 Bioactivity Elution Profiles

To enable quantitative monitoring of the amount of eluted molecule, F6 was tagged with a fluorescent marker, Cy5. The concentration of released F6 was then measured by means of a plate reader working in a fluorometric mode with the excitation wavelength at 675 nm and the emission wavelength at 690 nm. The calibration curve (***Figure 1***) was plotted for the fluorescence versus F6 concentration (where y is the fluorescence and x is the concentration in µg ml^−1^). A linear relationship was observed between 0.5 µg ml^−1^ and 500 µg ml^−1^ satisfying the equation y = 1.35 x + 2.56 (R^2^ = 0.9994).

**Figure 1.**
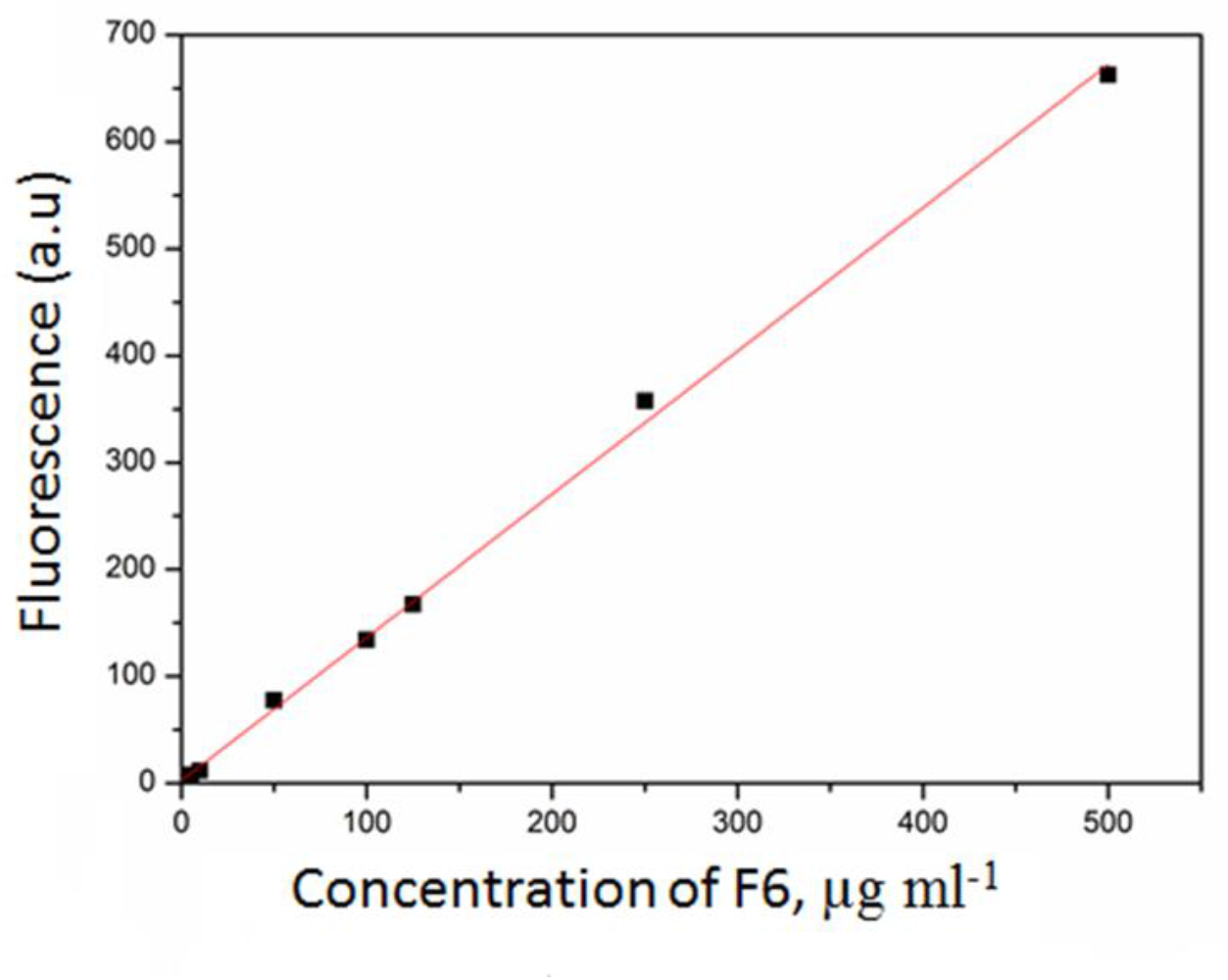
Calibration curve showing the linear relationship of the fluorescence heparan sulphate mimetic-F6-Cy5 tagged versus its concentration. A linear relationship is observed between 0.5 µg ml^−1^ and 500 µg ml^−1^ satisfying the equation y = 1.35 x + 2.56 (R2 = 0.9994).

##### - Passive Release Model

F6 elution in a spontaneous, passive mode was studied for the period of 21 days with the concentration measurements performed on a daily basis. To simulate *in vivo* conditions, PEDOT:PTS:F6 coated electrodes were immersed in phosphate-buffered saline solution (PBS 1X) solution, placed on a shaker and kept at the temperature of 37° C. Two 100 µl samples of the supernatant were taken from each sample and analysed by a plate reader working in fluorometric mode. Each time, 200 µl of a fresh PBS 1X solution was added to the sample to keep the total amount of supernatant at a constant level (2 ml). The continual addition of a fresh solution enabled the sink conditions to be maintained, i.e. prevented the saturation of the solution with F6.

##### - Active Release Model

The active release of F6 was triggered electrically through a biphasic potential pulse, consisting of a 5-second application of a reduction potential (−0.5 V) followed by a 5-second application of an oxidative potential (+0.5 V). The active release was studied for a period of 21 days with the concentration measurements performed on a daily basis. The electrical trigger consisting of 50 stimulation cycles was applied every daily. Two 100 µl samples of the supernatant were taken from each sample and analysed by a plate reader working in a fluorometric mode. Each time, 200 µl of a fresh PBS 1X solution was added to the sample to keep the total amount of supernatant constant (1 ml).

### 2.1.5 Cell Culture

Primary cultures of ventral mesencephalic neurons (VM) were obtained from the mesencephalon of embryonic Sprague–Dawley rats according to methods previously described by Vallejo-Giraldo et al. [5]. Briefly, the ventral mesencephalon were dissected from embryonic fourteen-day rat brains and then mechanically dissociated with a pipette, until the tissue was dispersed. Cells were grown in a humidified atmosphere of 5% CO_2_ at 37°C and culture in media (Dulbecco’s modified Eagle’s medium/F12, 33 mM D-glucose, 1 % L-glutamine, 1% PS, 1 % FCS, supplemented with 2 % B27). Controls and experimental groups were cultured for three, seven and ten days in six well culture plates and sterilised in 70% ethanol for two hours, and subsequently washed repeatedly with HBSS and/or molecular biology level water (Sigma). Prior to plating, samples and controls were coated with poly-lysine (PLL) (Sigma). They were then rinsed three times with molecular biology level water and left to dry overnight. 50,000 cells cm^−2^ were plated on each electrode, and then 3 ml of the culture medium was added to each well and half of the volume was replaced with fresh media every two days for a period of ten days.

For the inflammatory control, primary VM cells were cultured on sterile Thermanox® Plastic Coverslips with 13 mm diameter (NUNC^TM^ brand products). 50,000 cells cm^−2^ were plated on each coverslip, and grown in a humidified atmosphere of 5% CO_2_ at 37°C and culture in media (Dulbecco’s modified Eagle’s medium/F12, 33 mM D-glucose, 1 % L-glutamine, 1% PS, 1 % FCS, supplemented with 2 % B27). 3 ml of the culture medium was added to each well and, after two days in culture, the cells were stimulated with IL-1β (10 ng mL^−1^) prepared in plating media to a final volume of 3 ml and half of the volume was changed every two days for a period of ten days.

### 2.1.6 Immunofluorescent Labelling

Indirect double-immunofluorescent labelling was performed to visualise neurons and astrocyte cell populations as described previously [70]. Briefly, VM cells on experimental and control substrates were fixed with 4% paraformaldehyde and 1% of sucrose for twenty minutes at room temperature at the time point. Once fixed, the samples were washed with PBS and permeabilised with buffered 0.5% Triton X-100 within a buffered isotonic solution (10.3 g sucrose, 0.292 g NaCl, 0.06 g MgCl_2_, 0.476 g HEPES buffer, 0.5 ml Triton X 100, in 100 ml water, pH 7.2) at 4°C for five minutes. Non-specific binding sites were blocked with 1% bovine serum albumin (BSA) in PBS at 37°C for 30 minutes and subsequently incubated for two hours with a 1:200 concentration anti-glial fibrillary acidic protein (GFAP) antibody produced in mouse (Sigma, 1:200) and 1:500 concentration anti-β-Tubulin III antibody produced in rabbit (Sigma, 1:500). Samples were washed three times with 0.05% Tween 20/PBS and then incubated for one hour in the secondary antibody Alexa Fluor® 488 goat anti-Mouse IgG / IgA / IgM (H+L) (Molecular probes 1:500) combined with the secondary antibody Alexa Fluor® 594 goat anti-Rabbit IgG (H+L) (Molecular probes, 1:500). Samples were washed with PBS (five minutes ×3) and mounted on microscope cover slides and counterstained with slowfade^R^ gold antifade reagent with DAPI for nuclear staining.

### 2.1.7 Microscopy and Image Analysis

After immunostaining, samples were viewed with an Olympus Fluoview 1000 Confocal Microscope at a fixed scan size of 1024 by 1024 at a ratio 1:1. Cell analysis was performed as described in [70]. At least twenty images at 60× magnification were taken at random from each experimental group and controls. Cell density was analysed by counting the total number of labelled nuclei corresponding to neurons and astrocytes in an area of 211.97 µm * 211.97 µm. Neurite length was quantified by analysing nine random fields of view of three different technical replicas from three different samples using established stereological methods [71]. The formula used was: neurite length = n*T *π/2, where n is the number of times neurites intersect grid lines and T = distance between gridlines (taking magnification into account) as described in [72].

### 2.1.8 Cytokine Inflammatory Panel

Cytokine multiplex assay was performed on primary VM cell mixed population supernatants collected at three, seven and tend days in culture grown on all experimental groups, controls and inflamed control. ELISA pro-inflammatory panel 2 (Rat) (Meso Scale Discovery, UK) cytokine (IL-6, IL-1β, TNF-α, IFN-Υ, KC/GRO, IL-4, IL-5, IL-13, IL-10) assays were performed according to the manufacturer’s instructions, using six replicas and without adjustments to the recommended standard curve, and sample dilutions. Briefly, 150 µL of blocker H was added on each well of ELISA pro-inflammatory plate and incubated at room temperature with fast shaking for one hour. In parallel, in a separately provided plate, an initial 1:2 dilution of samples and culture media was prepared and put under shaking for fifteen minutes. The ELISA pro-inflammatory plate was then washed three times with at least 150 µL of wash buffer (PBS 1X with 0.05% Tween20) and subsequently, the diluted sample mixed from the additional plate provided was transferred to the ELISA pro-inflammatory plate with the addition of 25 µl of diluent 40. Incubation at room temperature with shaking for two hours was then carried out. The ELISA pro-inflammatory plate was further washed three times with PBS-Tween20, and 25 µL of 1X detection antibody solution was added in each well and incubated at room temperature with shaking for two hours. The ELISA pro-inflammatory plate was washed three times and further added 150 µL of 2X read buffer T in each well for plate reading using the QuickPlex SQ 120 multiplexing instrument from MSD.

### 2.1.9 Glycosaminoglycans Competition Assay Towards FGF-2 and VEGF Binding

F6 molecule [60], Heparin, HS and polymeric dextran (Sigma-Aldrich) binding to basic fibroblast growth factor (FGF-2, R&D systems) and Vascular Endothelial Growth Factor 165 (VEGF165, Promokine) were evaluated by an ELISA based competition assay as previous described [73]. 96 well ELISA plates were coated with a 2 µg mL^−1^ heparin-bovine serum albumin (BSA) conjugate solution prepared as previously described in [74]. Briefly, after washing 0.05% Tween 20/PBS, wells were saturated with 3% BSA in PBS. Then, FGF-2 (10 ng mL^−1^) and F6 molecule, heparin, HS or dextran, (in a concentration dependent manner; 0.001, 0.01, 0.1 1, 10 and 100 µg mL^−1^) were simultaneously added to wells. After one hour incubation at room temperature, wells were washed and the growth factors remained bond to the heparin conjugates were detected by incubation with the corresponding antibody (R&D systems) followed by a peroxidase-labeled secondary antibody (Jackson ImmunoResearch). Peroxidase activity measurement was performed with the 3,3’,5,5’-tetramethylbenzidine (TMB) substrate of the Peroxidase Activity Detection Kit (Pierce^TM^) following fabricant indications. FGF-2 and VEGF bindings to the heparin-BSA conjugate were considered as reference binding (100%).

### 2.1.10 Protein Antibody Microarray and Lectin Array

#### 2.1.10.1 Protein Extraction of Primary VM cell Mixed Population (VM) Cells and Labelling

For protein extraction and collection, samples were carefully placed on ice and the cells were washed twice with ice-cold PBS, pH 7.4. After the complete aspiration of ice-cold PBS, 100 µl of cold RIPA lysis buffer (Sigma) supplemented with 1% of protease inhibitors (Life Science-Roche) and 1% of phosphatase inhibitors cocktails I & III (Sigma) was added. Thereafter, the adherent cells were scraped off from each of the experimental groups and controls using Corning® cell scrapers and then the cell suspension was gently transferred into microcentrifuge tubes and placed on ice. Microcentrifuge tubes were centrifuged for fifteen minutes at 14,000 rpm at 4 °C, and afterwards gently removed from the centrifuge and placed on ice. Without disturbing the pellet, the supernatant was carefully aspirated and placed in a fresh clean tube, kept on ice, and stored at −80 °C to be quantified by bicinchoninic acid (BCA) protein assay kit [75] according to manufacturer’s instructions using a bovine serum albumin standard curve (ThermoFisher Scientific). The pellet was discarded.

Protein supernatants were then labelled with Alexa Fluor® 555 carboxylic acid succinimidyl ester (AF555) (ThermoFisher Scientific) essentially as previously described [76]. All processes were carried out in the dark. In brief, approximately 0.5 mg of each sample by protein concentration was incubated with approximately 50 µg of AF555 in 250 mM sodium bicarbonate, pH 8.2 for two hours at room temperature in the dark. Excess label was removed and buffer was exchanged with PBS, pH 7.4 by centrifugation through 3 kDa molecular weight cut off centrifugal filters (Amicon Ultra®, Millipore, Dublin, Ireland). Absorbance at 555 and 280 nm was measured for labelled samples and calculations were performed according to manufacturer’s instructions using an arbitrary extinction coefficient of 100,000 and molecular mass of 100,000 to enable quantification of relative protein concentration and label substitution efficiency. Labelled samples were stored at 4 °C protected from light.

#### 2.1.10.2 Construction of Antibody and Lectin Hybrid Microarray

All commercial antibodies (***Table 1***) were buffer exchanged in to PBS and quantified by BCA assay. A panel of 15 antibodies and 26 pure unlabelled lectins (***Table 2***) were printed in PBS, pH 7.4 on Nexterion® H amine reactive, hydrogel coated glass slides (Schott AG, Mainz, Germany) using a SciFLEXARRAYER S3 piezoelectric printer (Scienion, Berlin, Germany) under constant humidity (62% +/− 2%) and a constant temperature of 20°C essentially as previously described [5, 76]. The lectins were printed at 0.5 mg mL^−1^ in PBS supplemented with 1 mM of their respective haptenic monosaccharides (***Table 2, Figure 2***) while antibodies were printed at concentrations between 0.1 – 1.0 mg mL^−1^ in PBS (***Table 1, Figure 2***). Antibodies and lectins were printed in replicates of six and each feature was printed using approximately 1 nL using an uncoated 90 µm glass nozzle with eight replicate subarrays printed per microarray slide. After printing, slides were incubated in a humidity chamber overnight at room temperature to facilitate complete conjugation. The slides were then blocked in 100 mM ethanolamine in 50 mM sodium borate, pH 8.0, for one hour at room temperature. Slides were washed in PBS with 0.05% Tween® 20 (PBS-T) three times for three minutes each wash, followed by one wash in PBS, dried by centrifugation (470 x g, five minutes) and then stored with desiccant at 4 °C until use.

**Table 1.** Summary of all the commercial antibodies used for the gliosis antibody microarray.

| Order | Probe | Conc<br>[mg mL<sup>-1</sup>] | Information | Company | Catalogue<br>Number |
| --- | --- | --- | --- | --- | --- |
| 1 | Anti-Glial Fibrillary<br>Acidic Protein (GFAP) | 0.25 | Mouse anti-rat<br>immunoglobulins,<br>monoclonal | Sigma® | G3893 |
| 2 | Nestin (Rat-401) | 0.5 | Mouse anti-rat<br>immunoglobulins,<br>monoclonal | SantaCruz® | sc-33677 |
| 3 | CD81 (H-121) | 1 | Rabbit anti-rat<br>immunoglobulins,<br>polyclonal | SantaCruz® | sc-9158 |
| 4 | Integrin alphaM<br>(OX42) | 1 | Mouse anti-rat<br>immunoglobulins,<br>monoclonal | SantaCruz® | sc-53086 |
| 5 | Anti Iba1 | 2.5 | Rabbit anti-rat<br>immunoglobulins,<br>polyclonal | WAKO® | 019-19741 |
| 6 | Paxillin antibody<br>[Y113] | 0.5 | Rabbit anti-rat<br>immunoglobulins,<br>monoclonal | Abcam | ab32084 |
| 7 | cleaved spectrin alpha<br>II (h1186) | 0.5 | Rabbit anti-rat<br>immunoglobulins,<br>polyclonal | SantaCruz® | sc-23464 |
| 8 | Anti-Smad3 antibody<br>[EP568Y] | 1 | Rabbit anti-rat<br>immunoglobulins,<br>monoclonal | Abcam | ab40854 |
| 9 | Anti-Chondroitin<br>Sulfate antibody | 0.1 | Mouse anti-rat<br>immunoglobulins,<br>monoclonal | Sigma® | C8035 |
| 10 | PBS |  | Phosphate buffered<br>saline, pH 7.4 |  |  |
| 11 | Anti-rat_0.5 | 0.5 | Rabbit anti-rat immunoglobulins, polyclonal | DAKO |  |
| 12 | Anti-sheep_0.5 | 0.5 | Rabbit anti-sheep immunoglobulins, polyclonal | DAKO |  |
| 13 | VR1 (H-150) | 1 | Rabbit anti-human immunoglobulins, polyclonal | SantaCruz® | sc-20813 |
| 14 | Anticorps L-type Ca++ CP α1C (H-280) | 0.5 | Rabbit anti-human immunoglobulins, polyclonal | SantaCruz® | sc-25686 |
| 15 | PIEZO1 Antibody (N-15) | 1 | Goat anti-human immunoglobulins, monoclonal | SantaCruz® | sc-164319 |
| 16 | PIEZO2 Antibody (G-20) | 1 | Rabbit anti-human immunoglobulins, polyclonal | SantaCruz® | sc-84763 |
| 17 | ANKTM1 (C-19) | 0.5 | Goat anti-human immunoglobulins, polyclonal | SantaCruz® | sc-32353 |
| 18 | TREK-1 Antibody (C-20) | 1 | Goat anti-human immunoglobulins, polyclonal | SantaCruz® | sc-11557 |
| 19 | Anti β –Tubulin III (TUBB3) | 0.1 | Rabbit anti-rat immunoglobulins, polyclonal | Sigma® | T2200 |
| 20 | Anti β –Tubulin III (TUBB3) | 0.25 | Rabbit anti-rat immunoglobulins, polyclonal | Sigma® | T2200 |
| 21 | Anti β –Tubulin III (TUBB3) | 0.5 | Rabbit anti-rat immunoglobulins, polyclonal | Sigma® | T2200 |
| 22 | Anti β –Tubulin III (TUBB3) | 0.75 | Rabbit anti-rat immunoglobulins, polyclonal | Sigma® | T2200 |
| 23 | Anti β –Tubulin III (TUBB3) | 1 | Rabbit anti-rat immunoglobulins, polyclonal | Sigma® | T2200 |

**Table 2.** Lectins, their sources, common names, binding specificities, print sugars and the manufacturer.

| Abbreviation | Source | Species | Common name | General binding specificity | Print sugar | Supplier |
| --- | --- | --- | --- | --- | --- | --- |
| AIA (Jacalin) | Plant | *Artocarpus integrifolia* | Jack fruit lectin | Gal, Gal-β-(1,3)-GalNAc (sialylation independent) | Gal | EY Labs |
| SJA | Plant | *Sophora japonica* | Pagoda tree lectin | β-GalNAc | Gal | EY Labs |
| DBA | Plant | *Dolichos biflorus* | Horse gram lectin | α-GalNAc | Gal | EY Labs |
| WFA | Plant | *Wisteria floribunda* | Japanese wisteria lectin | GalNAc/sulfated GalNAc | Gal | EY Labs |
| HPA | Animal | *Helix pomatia* | Edible snail lectin | α-GalNAc | Gal | EY Labs |
| ACA | Plant | *Amaranthus caudatus* | Amaranthin | Sialylated/Gal-β-(1,3)-GalNAc | Lac | Vector Labs |
| GS-II | Plant | *Griffonia simplicifolia* | Griffonia/Bandeiraea lectin-II | GlcNAc | GlcNAc | EY Labs |
| sWGA | Plant | *Triticum vulgaris* | Succinyl WGA | GlcNAc | GlcNAc | EY Labs |
| NPA | Plant | *Narcissus pseudonarcissus* | Daffodil lectin | α-(1,6)-Man | Man | EY Labs |
| GNA | Plant | *Galanthus nivalis* | Snowdrop lectin | Man-α-(1,3)-linked | Man | EY Labs |
| Con A | Plant | *Canavalia ensiformis* | Jack bean lectin | α-Man / α-Glc | Man | EY Labs |
| Lch-B | Plant | *Lens culinaris* | Lentil isolectin B | α-Man, core fucosylated, agalactosylated biantennary N-glycans | Man | EY Labs |
| Lch-A | Plant | *Lens culinaris* | Lentil isolectin A | α-Man / α-Fuc | Man | EY Labs |
| PSA | Plant | *Pisum sativum* | Pea lectin | Branched α-Man, core fucosylated trimannosyl N-glycans | Man | EY Labs |
| WGA | Plant | *Triticum vulgaris* | Wheat germ agglutinin | NeuAc/GlcNAc | GlcNAc | EY Labs |
| MAA | Plant | *Maackia amurensis* | Maackia agglutinin | Sialic acid-α-(2,3)-linked | Lac | EY Labs |
| SNA-I | Plant | *Sambucus nigra* | Sambucus lectin-I | Sialic acid-α-(2,6)-linked | Lac | EY Labs |
| RCA-I/120 | Plant | *Ricinus communis* | Castor bean lectin-I | Gal-β-(1,4)-GlcNAc | Gal | Vector Labs |
| CAA | Plant | *Caragana arborescens* | Pea tree lectin | Complex oligosaccharides | Lac | EY Labs |
| ECA | Plant | *Erythrina cristagalli* | Cockscomb/coral tree lectin | Gal-β-(1,4)-GlcNAc oligomers, LacNAc | Lac | EY Labs |
| AAL | Fungi | *Aleuria aurantia* | Orange peel fungus lectin | Fuc-α-(1,6)-linked > Fuc-α-(1,3)- and -α-(1,2)-linked | Fuc | EY Labs |
| LTA | Plant | *Lotus tetragonolobus* | Lotus lectin | Fuc-α-(1,3)-linked | Fuc | EY Labs |
| UEA-I | Plant | *Ulex europaeus* | Gorse lectin-I | Fuc-α-(1,2)-linked | Fuc | EY Labs |
| MPA | Plant | *Maclura pomifera* | Osage orange lectin | Terminal Gal-β-(1,3)-GalNAc > GalNAc-α-(1,6)-Gal | Gal | EY Labs |
| VRA | Plant | *Vigna radiata* | Mung bean lectin | Terminal α-Gal | Gal | EY Labs |
| MOA | Fungi | *Marasmius*<br>*oreades* | Fairy ring<br>mushroom<br>lectin | Terminal α-Gal | Gal | EY Labs |

**Figure 2.**
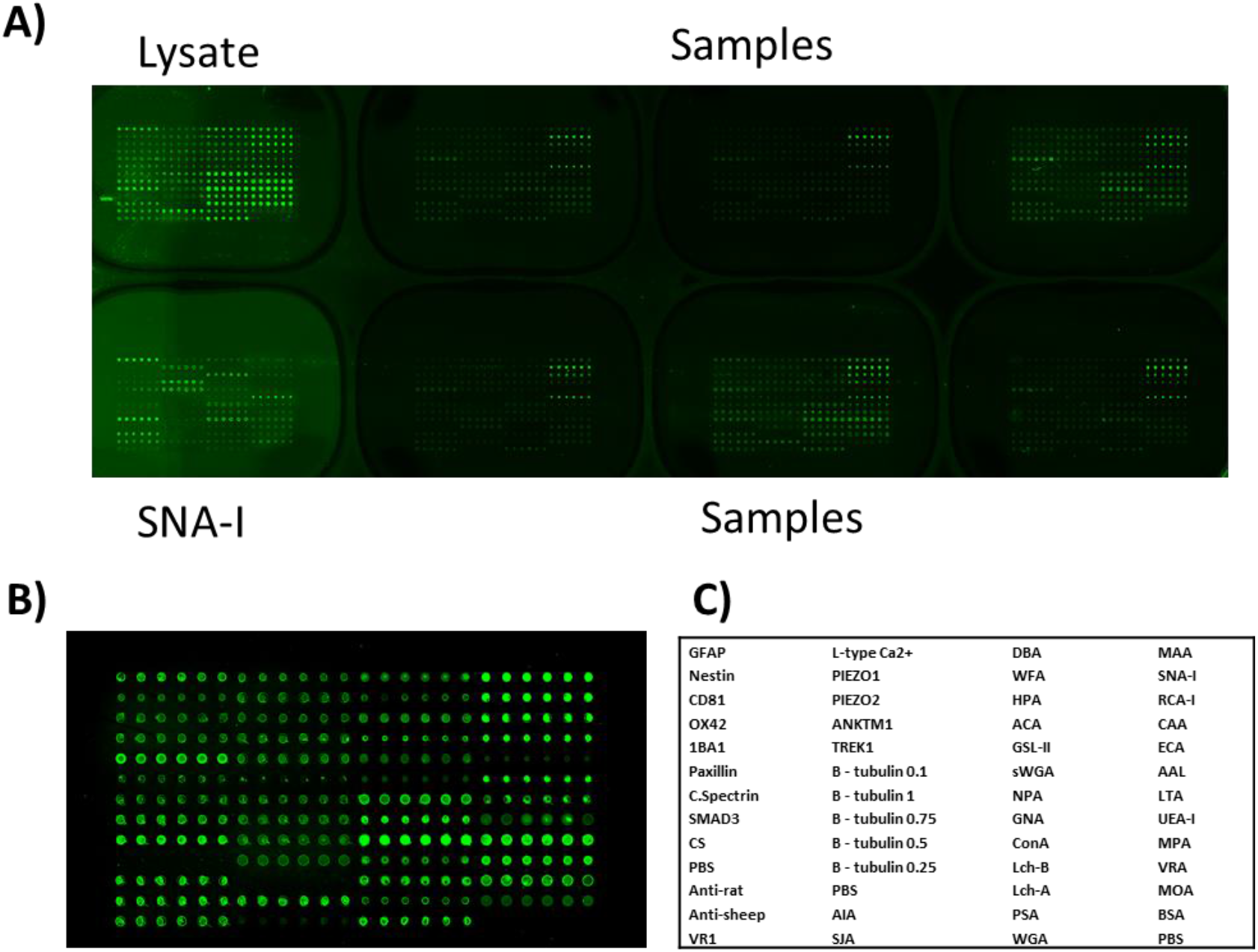
Images from a representative scanned microarray slide after incubation and washing. A) Incubations were carried out on three replicate microarray slides. Alexa Fluor® 555 labelled healthy rat brain lysate (5 mg mL-1) and SNA-I lectin (10 mg mL-1) were used as controls to confirm retained antibody performance and printing, respectively, and were incubated in two separate subarrays on every slide. B) Individual subarray from a representative scanned microarray slide after incubation with healthy rat brain lysate and subsequent washing. The subarrays were printed with antibodies and lectins in replicates of six following the order detailed in C).

#### 2.1.10.3 Profiling Samples on Hybrid Microarrays, Data Extraction and Analysis

All incubations were carried out under dark conditions. Microarray slides were incubated as previously described [5, 76]. Initially, one labelled sample was titrated (2.5 - 15 µg mL^−1^) to determine optimal signal to noise ratio and all samples were subsequently incubated for one hour at 23 °C at 5 µg mL^−1^ in Tris-buffered saline (TBS; 20 mM Tris-HCl, 100 mM NaCl, 1 mM CaCl2, 1 mM MgCl_2_, pH 7.2) with 0.05% Tween® 20 (TBS-T). All microarray experiments were carried out using three replicate slides. AF555-labelled healthy rat brain lysate (5 µg mL-1) and tetramethylrhodamine-(TRITC)-labelled SNA-I lectin (10 µg mL^−1^) were incubated in two separate subarrays on every slide to confirm retained antibody performance and printing, respectively (***Figure 2***). After incubation, slides washed three times in TBS-T for three minutes per wash, once in TBS and then centrifuged dry as above. Dried slides were scanned immediately on an Agilent G2505 microarray scanner using the Cy3 channel (532 nm excitation, 90% PMT, 5 µm resolution) and intensity data was saved as a .tif file. The antibodies and lectins conjugated to the microarray surface were verified to remain active for at least one month after printing and all incubations were carried out within that timeframe.

Data extraction from .tif files was performed essentially as previously described [5, 76]. Data were normalised to the mean of three replicate microarray slides (subarray by subarray using subarray total intensity, N= 3, 18 data points). Unsupervised hierarchical clustering of normalised data was performed using Hierarchical Clustering Explorer v3.0 (http://www.cs.umd.edu/hcil/hce/hce3.html) using the parameters no pre-filtering, complete linkage, and Euclidean distance. Statistical analysis was performed as detailed in the section 2.3.11.

### 2.1.11 Statistical Analysis

All data presented here was confirmed using at least three replicates for each of the test groups and control groups. The results are expressed as the mean of the values ± standard error of the mean. One-way ANOVA followed by a Bonferroni test were performed to determine the statistical significance (p<0.05), unless otherwise stated.

### 2.2 Results and Discussion

The use of GAGs as dopants within an electrodeposited polymer not only endows the resulting polymer with bioactivity, but as charged biomolecules, also act as counterions in the electrochemical polymerisation process [28, 36]. Experimentally, this study employed a PEDOT derivative, modified through the incorporation of the HM - F6, a polysulphated molecule which acted as a biological co-dopant for the formulation of glycan functionalised neural interfaces [52, 60, 77, 78]. F6 was originally described by Papy-Garcia et al. as a functionalised dextran derivative with resistance to glycanase degradation [44]. F6 together with the conventional sulphonate anion PTS, was incorporated in the matrix of PEDOT through a co-doping electropolymerisation process. The chemical structure of the F6 molecule and a proposed pictorial representation of PEDOT:PTS:F6 functionalisation on the physico-electrical and biological properties though F6 interaction are shown in ***Figure 3***.

**Figure 3.**
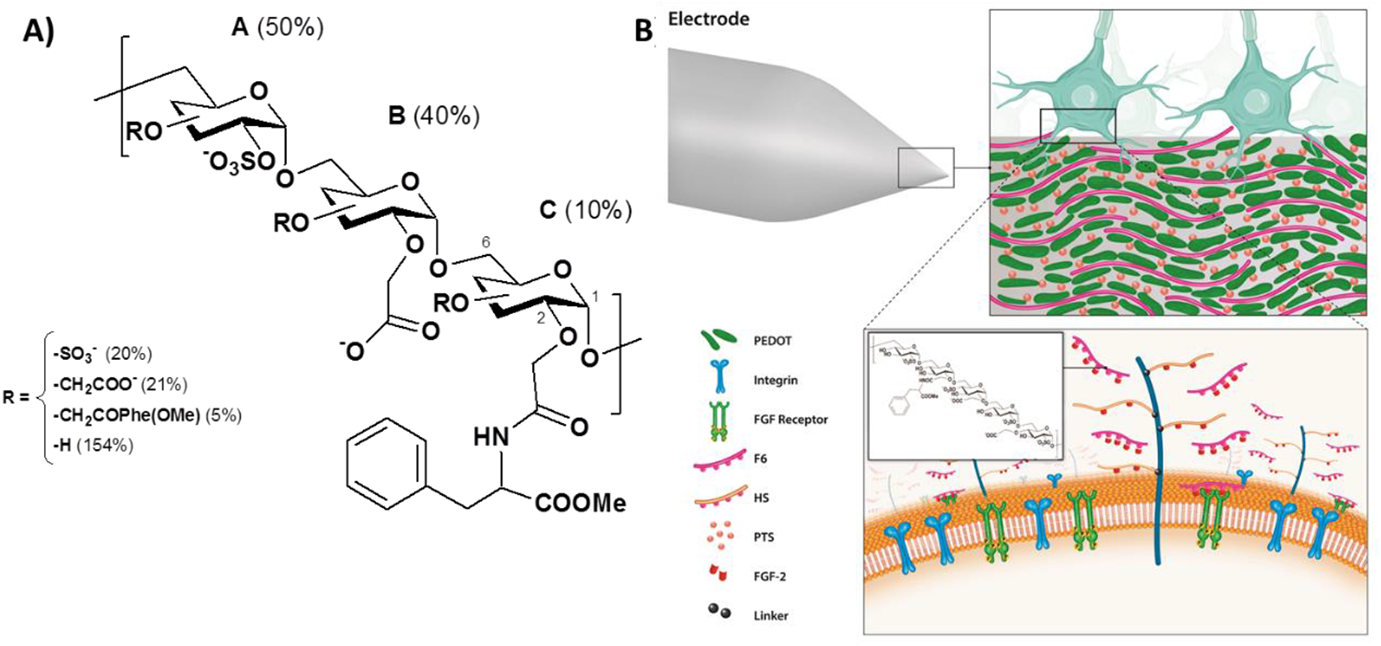
Biochemical functionalisation of the neuroelectrode surface with PEDOT:PTS:F6 thin films. A) F6 is a dextran T5 (Mw 5,000 Daltons) derivative containing sulphate, carboxymethyl, and phenylalanine methyl ester groups. For an easy representation, three differently substituted glucosidic units (A, B, and C) are depicted and arranged in an arbitrary combination. The relative content of each unit was calculated according to the nature of the group substituted at position 2 of each unit (C2). R represents the proportion of each substituted group at C3 and C4, which total substitution corresponds to 200%. F6 was synthesised and analysed as in [44]. B) Pictorial representation of PEDOT:PTS:F6 functionalised electrodes from the electrophysical arrangement to the cell level interaction with growth factors binding, i.e FGF-2.

Initially, an optimisation process to assess the therapeutic concentration of F6 was conducted with concentrations of 1, 10, 50, 100 and 1000 µg ml^−1^ incorporated into cell culture media over a period of ten days in culture. Cell viability effects using primary ventral mesencephalic mixed neural cell population grown in vitro are presented in ***Figure 4***. Live/dead analysis indicated maintained cell viability when F6 was present in the cell culture media at concentrations below 50 µg ml^−1^, a range that was replicated in F6 functionalised PEDOT:PTS films electrodeposited from an electrolyte/EDOT solution containing 1000 µg ml^−1^ of F6 (***Table 3***).

**Figure 4.**
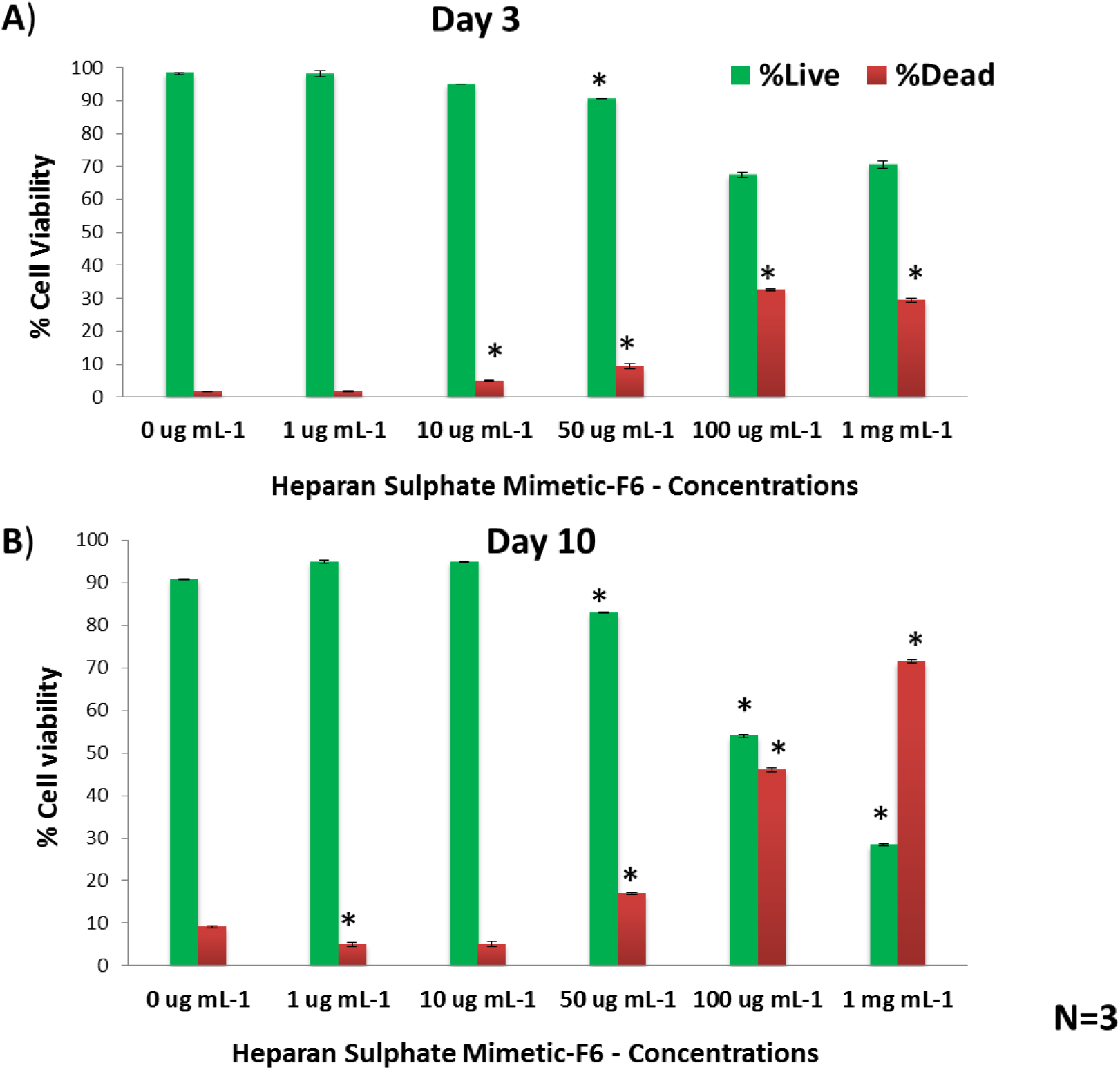
Cytocompatibility effects of heparan sulphate mimetic-F6 – concentrations in solution. Live and dead assay was carried out using primary ventral mesencephalic (VM) mixed neural cell population under the presence of bulk F6 at 1, 10, 50, 100 and 1000 ug ml^−1^. An early time of three days in culture A) and late time of ten days B) were evaluated. A maintained cell viability was achieved when bulk F6 was present in the cell culture media at concentrations below 50 ug ml^−1^. Results are ± STD. ★ = p< 0.05.

**Table 3.** Passive mode concentration profile of heparan sulphate mimetic-F6 released from PEDOT:PTS:F6 coated electrodes at a concentration of 1, 10, 50, 100 and 1000 ug ml^−1^ for a period of 21 days. Results are ± STD. N=3

| | | PEDOT:PTS:F6<br>at<br>1000 $\mu\text{g ml}^{-1}$ | PEDOT:PTS:F6<br>at<br>100 $\mu\text{g ml}^{-1}$ | PEDOT:PTS:F6<br>at<br>50 $\mu\text{g ml}^{-1}$ | PEDOT:PTS:F6<br>at<br>10 $\mu\text{g ml}^{-1}$ | PEDOT:PTS:F6<br>at<br>1 $\mu\text{g ml}^{-1}$ |
| --- | --- | --- | --- | --- | --- | --- |
| Concentration of Heparan Sulphate Mimetic F6<br>Released from PEDOT:PTS polymeric Matrix<br>$\mu\text{g ml}^{-1}\text{cm}^{-2}$ | 3 days | 5.6 ± 1.8 | 2.1 ± 0.7 | 2.5 ± 1.1 | 1.8 ± 0.4 | 1.4 ± 0.4 |
| | 7 days | 8.0 ± 3.5 | 3.9 ± 1.1 | 3.5 ± 1.4 | 3.5 ± 0.4 | 1.8 ± 0.7 |
| | 10 days | 10.2 ± 3.2 | 5.6 ± 2.5 | 6.0 ± 1.4 | 4.9 ± 0.7 | 3.2 ± 1.4 |
| | Total<br>amount<br>of F6<br>released<br>over 21<br>days | 11.7 ± 2.8 | 5.6 ± 3.5 | 6.0 ± 1.8 | 6.7 ± 0.7 | 5.3 ± 1.8 |

Additionally, PEDOT:PTS films electrodeposited from an electrolyte/EDOT solution containing 1000 µg ml^−1^ of F6 were found to possess the highest mass of immobilised F6 within the polymer matrix as assessed via Electrochemical Quartz Crystal Microbalance (EQCM)

***Figure 5***. Taken together, the optimal F6 electrolyte/EDOT solution concentration selected for subsequent analysis was 1000 µg ml^−1^, denoted hereon as “PEDOT:PTS:F6” in this work. PEDOT:PTS doped with a non-sulphated polymeric dextran was also employed as a control group to understand further the effect of sulphation pattern provided by F6. This was incorporated during the electrodeposition process of EDOT at the same concentration of F6, 1000 µg ml^−1^.

**Figure 5.**
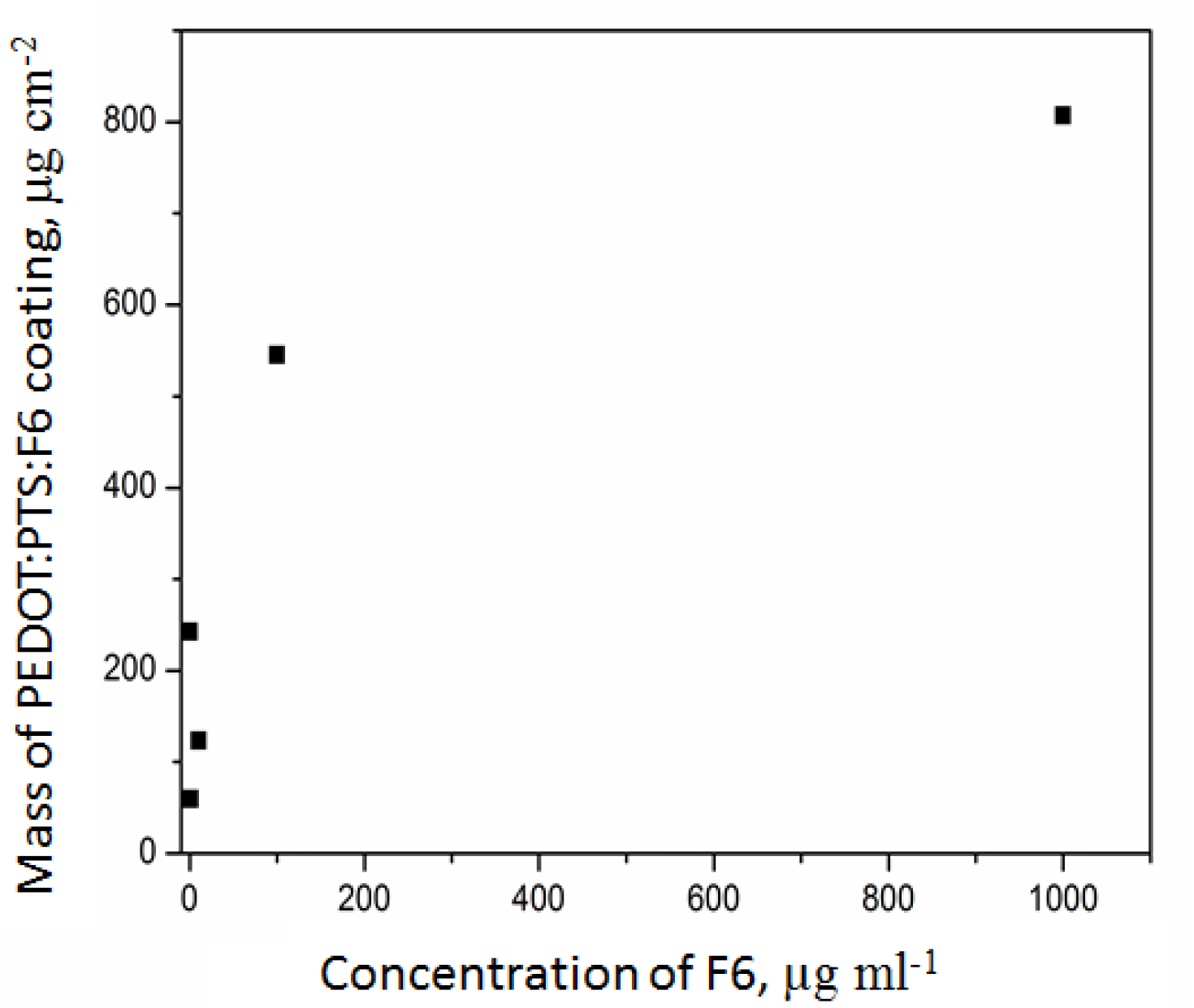
Electrochemical quartz crystal microbalance (EQCM) measurements of the masses recorded of PEDOT:PTS:F6 coatings at 1, 10, 50, 100 and 1000 µg ml^−1^. PEDOT:PTS:F6 at 1000 µg ml^−1^ of heparan sulphate mimetic-F6 was found to possess the highest mass immobilised of F6 within the polymer matrix with a mass recorded of 807.2 µg cm^−2^.

#### 2.2.1 Physical and Chemical Characterisation of Electrodeposited PEDOT:PTS:F6 Films

Electrodeposition of PEDOT:PTS:F6 onto concentric platinum iridium (PtIr) microelectrodes was confined to the conducting surfaces and did not result in an increase of electrode area at the macroscale. Representative SEM micrographs of the bare PtIr A) and the PEDOT:PTS:F6 coated microelectrodes B) are indicated in ***Figure 6***. The experimental values of the roughness and thickness profiles for the pristine PEDOT:PTS coatings and the functionalised PEDOT:PTS:F6 coatings at different F6 concentrations are detailed in ***Table 4***. The average roughness, R_a_ of electrodeposited PEDOT:PTS:F6 films was observed to decrease in a non-linear concentration dependent manner, but all electrodeposited PEDOT:PTS:F6 films possessed higher R_a_ values relative to pristine PEDOT:PTS films, a phenomenon reported previously with biological doping of PEDOT derivatives [79]. Relative to pristine PEDOT:PTS coatings, electrodeposited PEDOT:PTS:F6 electrode coatings possessed a denser and more uniform surface topography with more island-like particles, whereas films formulated with F6 concentrations < 1000 µg ml^−1^ possessed a less uniform and more porous surface structure (***Figure 7*A**). These observations taken in conjunction with FT-MIR spectra data indicate that with an increase in the concentration of F6 within the PEDOT:PTS polymer (***Figure 7*B**), the IR bands become noticeably sharper, owing to an increase in rigidity by restriction of conformational freedom [80].

**Figure 6.**
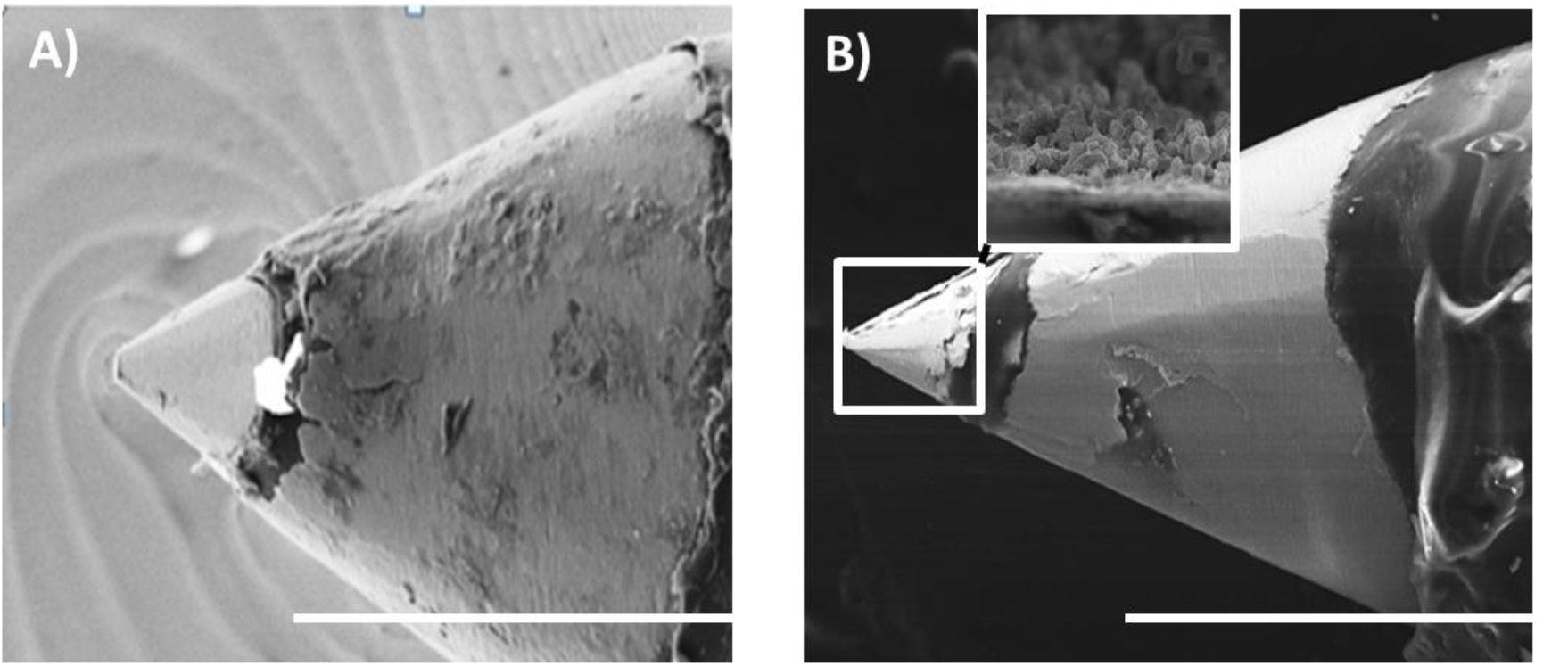
Scanning electron micrographs (SEM) of the bare platinum/iridium (PtIr) microelectrode A) and PEDOT:PTS:F6 coated microelectrode B) are shown, respectively. Scale bar = 200 µm. In the inset, the PEDOT:PTS:F6 coating is detailed, showing the nodular formation of electrodeposited coatings.

**Table 4.** Values of experimental mean surface roughness (R_a_) and mean thickness measurements of pristine PEDOT:PTS coated microelectrodes and PEDOT:PTS:F6 coated microelectrodes at 1, 10, 50, 100 and 1000 ug ml^−1^ of heparan sulphate mimetic-F6. The data represent the mean of 15 measurements from three different replicas. Results are ± SD, N=3.

| Electrodes | Average Roughness $R_a$<br>[nm] | Thickness<br>[nm] |
| --- | --- | --- |
| Pristine PEDOT:PTS | $85.16 \pm 16.94$ | $1210.01 \pm 0.08$ |
| PEDOT:PTS :F6 1 $\mu\text{g ml}^{-1}$ | $160.59 \pm 11.92$ | $1374.02 \pm 0.37$ |
| PEDOT:PTS :F6 10 $\mu\text{g ml}^{-1}$ | $167.00 \pm 25.12$ | $1619.00 \pm 0.41$ |
| PEDOT:PTS :F6 50 $\mu\text{g ml}^{-1}$ | $181.40 \pm 28.73$ | $784.80 \pm 0.24$ |
| PEDOT:PTS :F6 100 $\mu\text{g ml}^{-1}$ | $150.35 \pm 9.65$ | $2167.30 \pm 0.31$ |
| PEDOT:PTS :F6 1000 $\mu\text{g ml}^{-1}$ | $104.72 \pm 6.64$ | $2404.22 \pm 0.53$ |

**Figure 7.**
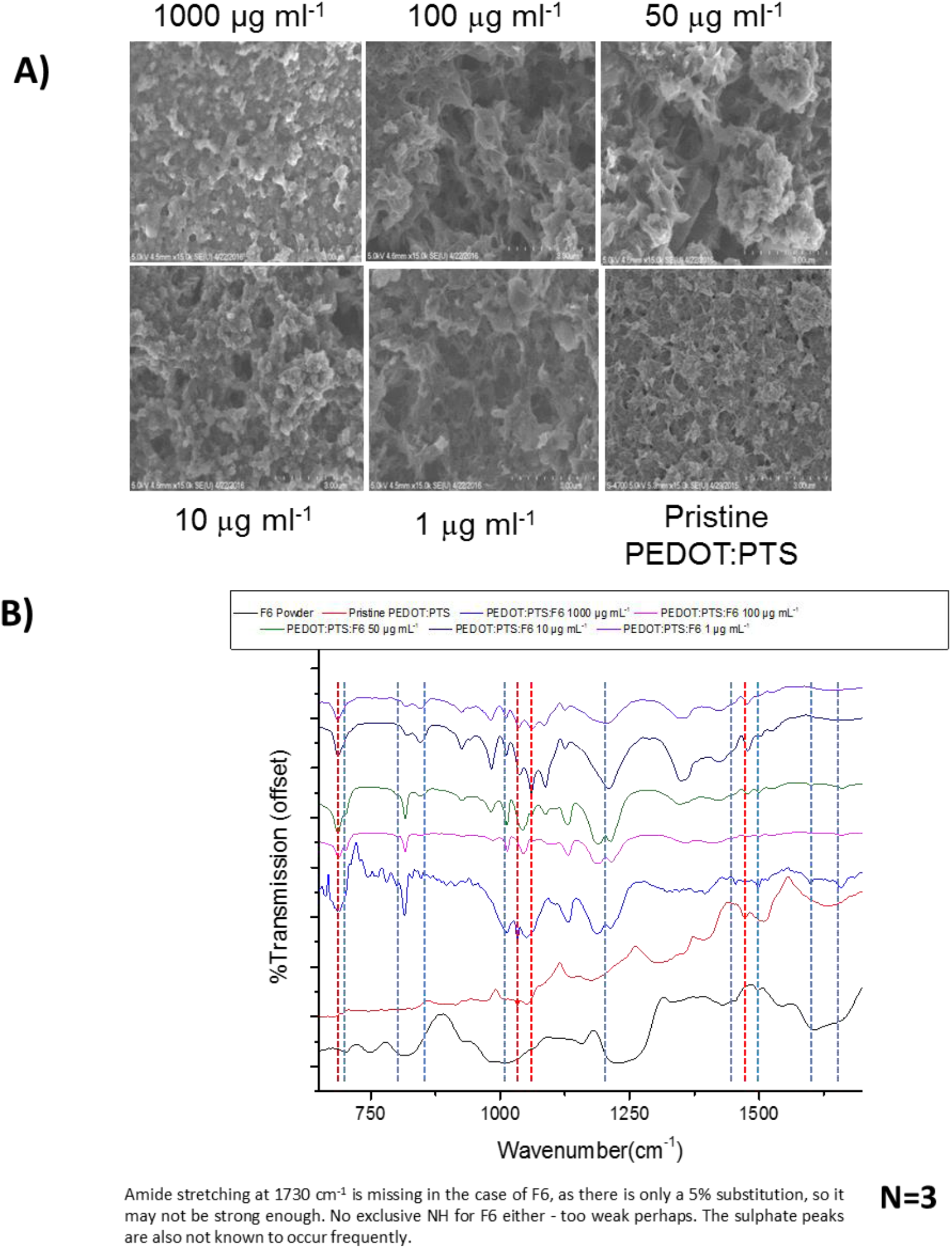
Morphological changes with different concentrations of F6. A) Scanning electron micrographs (SEM) of pristine PEDOT:PTS coated microelectrodes and the PEDOT:PTS:F6 coated microelectrodes at 1, 10, 50, 100 and 1000 µg ml^−1^ of heparan sulphate mimetic-F6. B) FT-MIR spectra of powdered F6 (black), pristine PEDOT:PTS coated microelectrodes (red) and PEDOT:PTS:F6 coated microelectrodes at 1, 10, 50, 100 and 1000 µg ml^−1^ of F6. Blue dotted lines show bands that belong to only F6, while red dotted lines show bands contributed by just PEDOT:PTS polymer. It can be seen that with an increase in the concentration of F6 in the PEDOT:PTS matrix, the bands become sharper suggesting a greater rigidity in the structure.

In order to assess F6 incorporation into electrodeposited PEDOT:PTS:F6 and pristine PEDOT:PTS films extensive chemical characterisation of coated microelectrodes was performed (***Figure 8*A-I**). Surface wettability of the electrodeposited films was assessed through contact angle analysis via an in-house developed goniometer. Pristine PEDOT:PTS coated microelectrodes were associated with a water contact angle of 15.92° ± 2.91, indicating the high hydrophilicity of the coatings [81]. Through the incorporation of F6, the water contact angle was observed to decrease to <5° accompanied by a rapid spreading of the water drop, indicating that the hydrophilicity of the electrodeposited films was significantly increased through doping with F6, potentially due to the presence of sulphate groups at the film surface, increasing polar interaction with the water droplet (***Figure 8*A**) [81]. Furthermore, the observed increase in PEDOT:PTS film roughness associated with F6 doping relative to pristine PEDOT:PTS coatings complements the observation of increased wettability of these coatings [82, 83].

**Figure 8.**
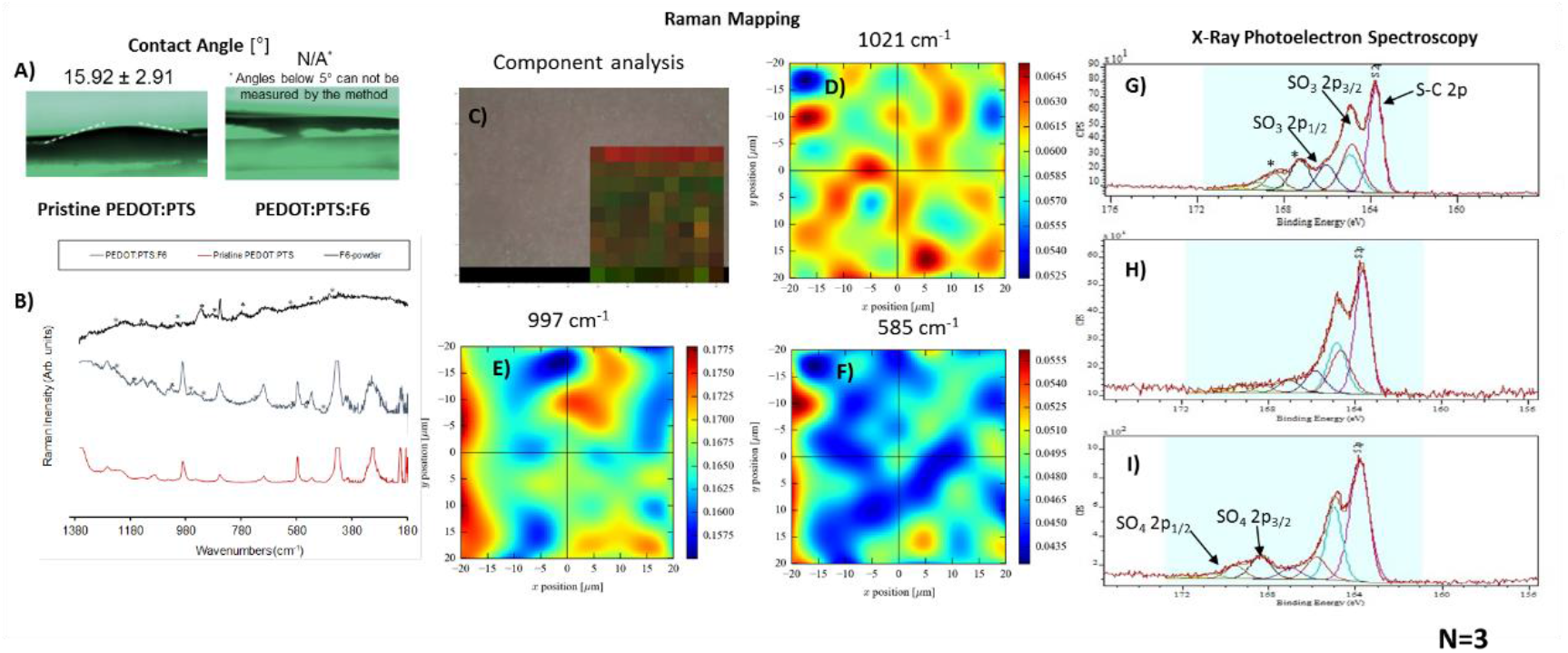
Chemical characterisation of the PEDOT:PTS:F6 coated microelectrode. A) Contact angle on pristine PEDOT:PTS and PEDOT:PTS:F6 coated microelectrodes, showing the decrease in a water contact angle suggesting an increase in hydrophilicity. B) Raman spectra of PEDOT:PTS microelectrodes, with and without incorporated F6, and powdered F6. Peaks assigned solely to F6 are indicated by an * on the pure F6 and of PEDOT:PTS:F6 coated microelectrodes spectra. (C-F) Raman mapping of pristine PEDOT:PTS and PEDOT:PTS:F6 coated microelectrodes. C) shows the component DLS analysis, where red pixels represent the F6 component and green pixels represent the PEDOT:PTS component. D) and E) show the intensity of the Raman bands at 1021 cm^−1^ and 997 cm^−1^, respectively, and indicate the presence of F6 evenly distributed on coated PEDOT:PTS microelectrodes. F) shows the band intensity at 585 cm^−1^ and indicates the presence of PEDOT:PTS in the coating. (G-I) X-ray photoelectron spectra of G) pristine PEDOT:PTS, H) PEDOT:PTS:Dextran, and I) PEDOT:PTS:F6 coated microelectrodes, showing the presence of sulphur in various chemical environments corresponding to S-C bonding (2p neutral state) and 2p_3/2_ and 2p_1/2_ states in both sulphonates and sulphates. The * in G) denotes the presence of possible impurities from the electrolyte.

Raman spectral analysis also indicated the presence of F6 in electrodeposited PEDOT:PTS:F6 coated microelectrodes ***Figure 8* B**). Peaks were observed at 997 cm^−1^ and 1021 cm^−1^ corresponding to C-C stretches and C-H deformations also observed in pristine F6. Peaks marked with * indicate that the microelectrode had measurable amounts of F6 incorporated into the PEDOT:PTS:F6 coating. The tentative assignments of peaks in the Raman spectra are shown in ***Table 5***. Peaks observed at 816 cm^−1^ and 848 cm^−1^, visible in both Raman and FT-MIR spectra (***Figure 7*B**), also confirm the presence of the F6 on the PEDOT:PTS:F6 coated microelectrodes. These bands correspond to E-E and E-A fragment γC-H deformations, respectively, in the dextran moiety [84]. Other peaks corresponding to F6 are indicated by blue dotted lines, and bands corresponding to the pristine PEDOT:PTS polymer are marked by red dotted lines in the IR spectra (***Figure 7*B**). A peak at 1180 cm^−1^ is masked by a board peak in spectra recorded from the two lower concentrations of F6, but emerges as a sharp peak as the F6 concentration increases. This band corresponds to C-O stretching [85]. Peaks at 928cm^−1^ in F6 (***Figure 7*B**) and 915 cm^−1^ in dextran (***Figure 9***) suggest the presence of the C1 gluocopyranasol conformation [84]. The presence of a peak at 800 cm^−1^ assigned to dextran suggests the presence of less than 5% alpha (1,3) glucoside bonds, making it highly linear [84] (***Figure 9***). The tentative assignments of peaks in the ATR-IR spectra are summarised in ***Table 6***.

**Figure 9.**
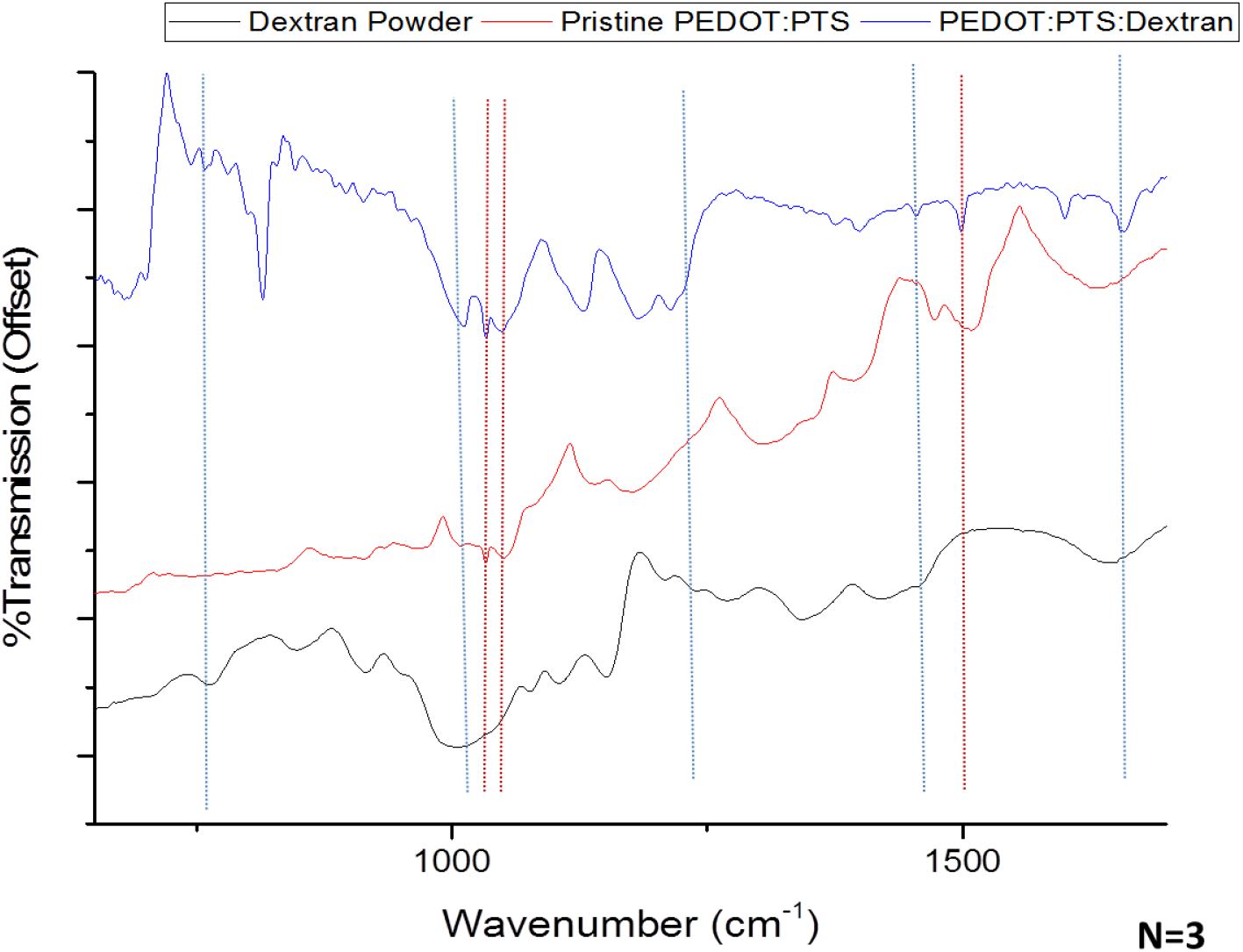
FT-MIR spectra of powdered dextran (black), pristine PEDOT:PTS coated microelectrodes (red) and PEDOT:PTS:Dextran coated microelectrodes. Blue dotted lines show bands that belong to only dextran, while red dotted lines show bands contributed just by PEDOT:PTS matrix.

**Table 5.**
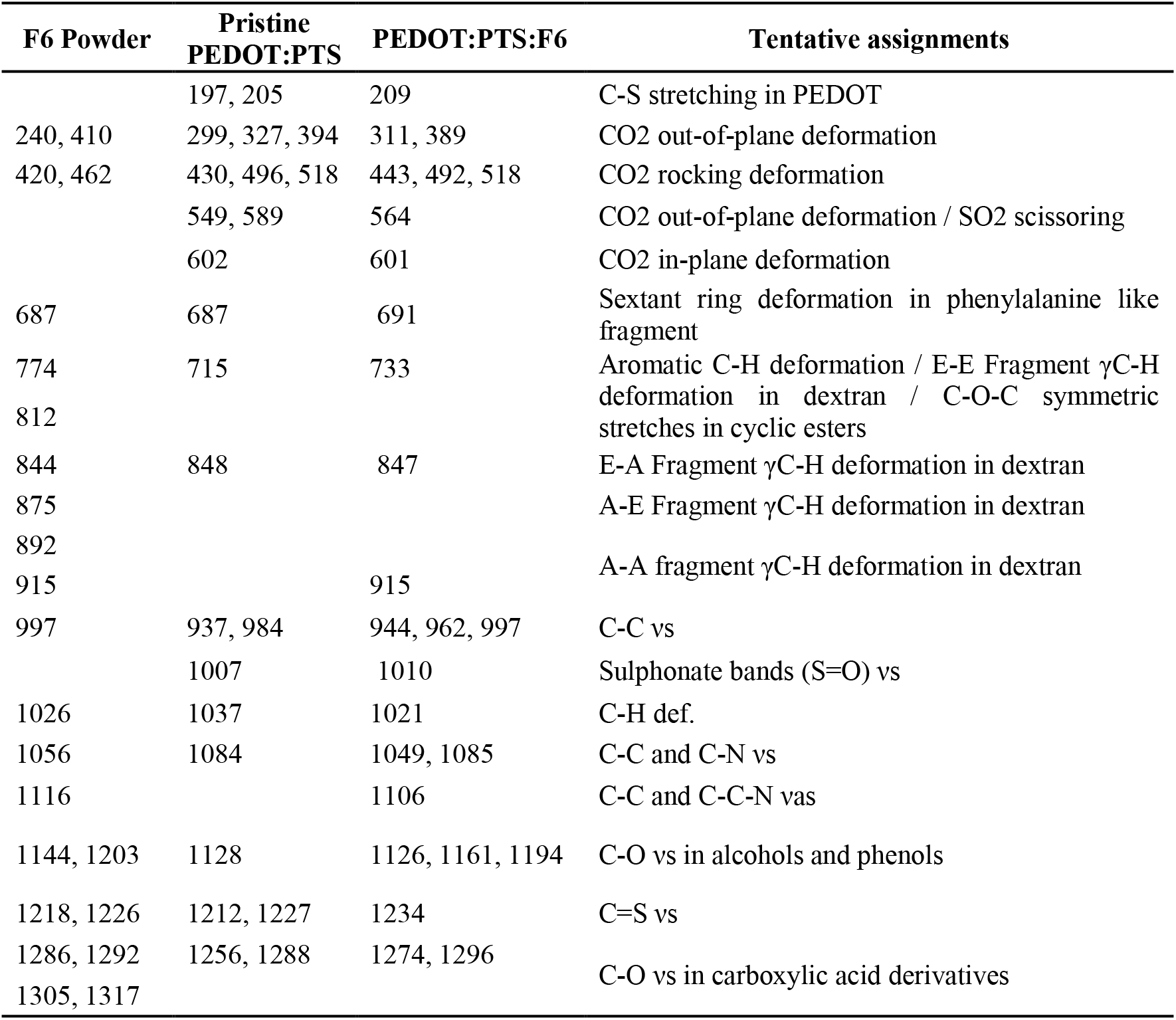
Tentative assignments of the peaks observed in the Raman spectra.

**Table 6.**
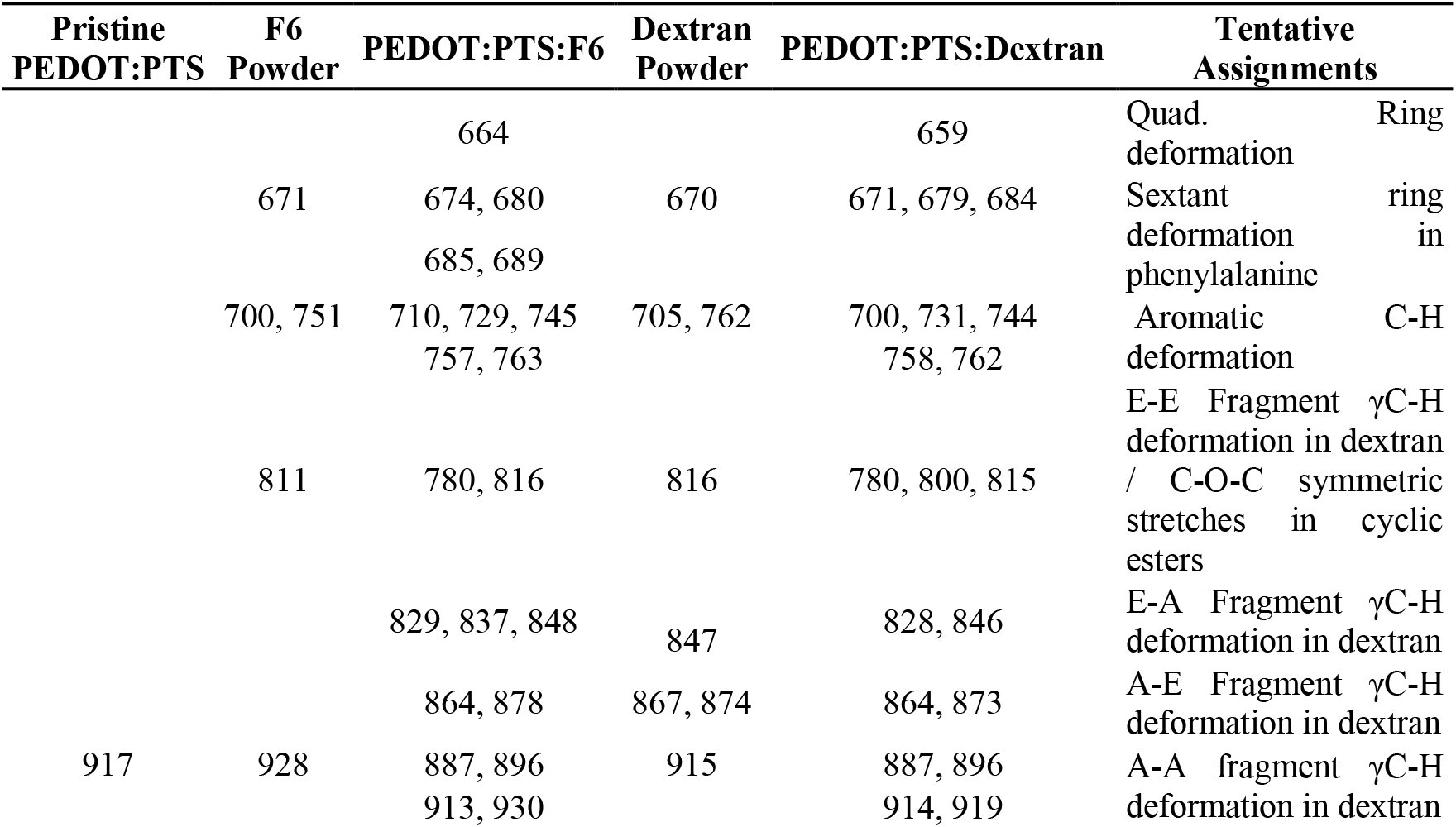
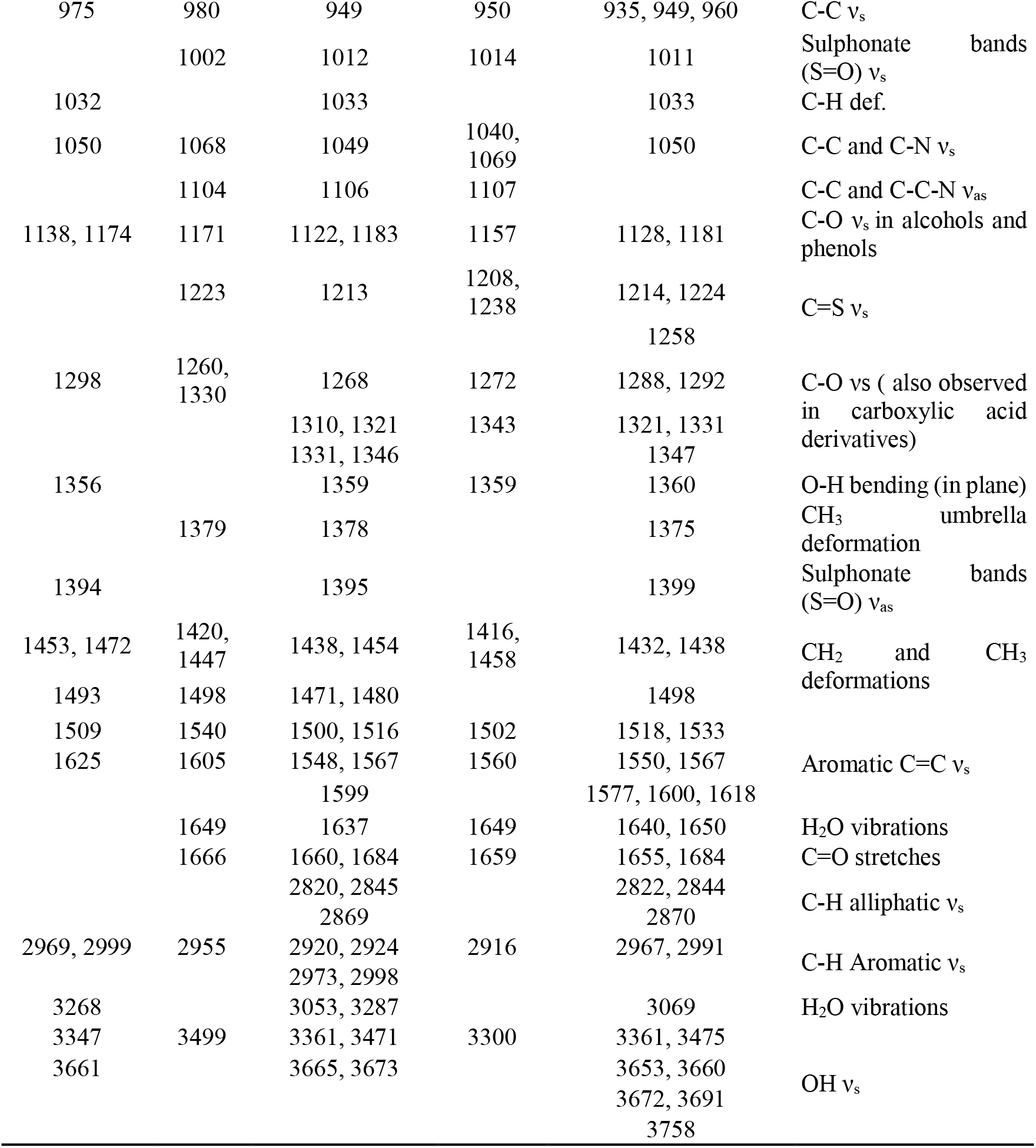
Tentative assignments of the peaks observed in the ATR-IR spectra.

Further, Raman mapping followed by component DLS analysis was performed to identify the F6 and the PEDOT:PTS components in electrodeposited films. The 585 cm^−1^ band corresponds to SO_2_ scissoring vibrations from PEDOT:PTS polymer, while bands at 997 cm^−1^ and 1021 cm^−1^ correspond to C-C stretches and C-H deformations observed in F6. ***Figure 8*C** shows a component analysis of these bands, where red pixels represent the F6 component and green pixels represent the PEDOT:PTS component, suggesting a uniform distribution of the F6 within the PEDOT:PTS:F6 coated microelectrodes. On comparing ***Figure 8*D** to ***Figure 8*E** and ***Figure 8*F**, a greater contribution from the C-H deformation in PEDOT:PTS:F6 films relative to the PEDOT:PTS films was observed. The highest-intensity contributions however were from the C-C stretches (***Figure 8*E**). These results successfully confirmed the presence of F6 on the surface of the PEDOT:PTS:F6 coated microelectrodes, with the increase in hydrophilicity observed from the contact-angle studies attributed to the sulphate groups from F6 being presented at the polymer surface.

Sulphate group vibrations are known to be weak and do not always appear in vibrational spectra, especially in the presence of other strongly absorbing groups [85, 86]. Therefore, to discern the chemical composition of PEDOT:PTS:F6 coating surface, X-ray photoelectron spectroscopy (XPS) analysis was conducted. ***Figure 8*(G-I)** show the surface analysis of pristine PEDOT:PTS, PEDOT:PTS:Dextran and the PEDOT:PTS:F6 coated microelectrodes, respectively. All coated electrodes were associated with XPS peaks corresponding to neutral S, arising from the S-C bonds in PEDOT (163.7 eV) (***Figure 8*G**) [87], and peaks arising from the 2p_3/2_ and 2p_1/2_ states of the sulphonates present in PTS (164.7 and 165.2 eV) (***Figure 8*H**) [88]. The presence of peaks arising from the 2p_3/2_ and 2p_1/2_ states of the sulphates from F6 (168.5 and 169.4 eV) were also observed (***Figure 8*I**) [89]. This peak is not present in the PEDOT:PTS:Dextran coated electrodes, while there appears to be some sulphate impurity in the pristine PEDOT:PTS coated electrodes, which could be from the electrolyte or formed during the electrodeposition process. The positions of C peaks and the relative compositions can be found in ***Figure 10***.

**Figure 10.**
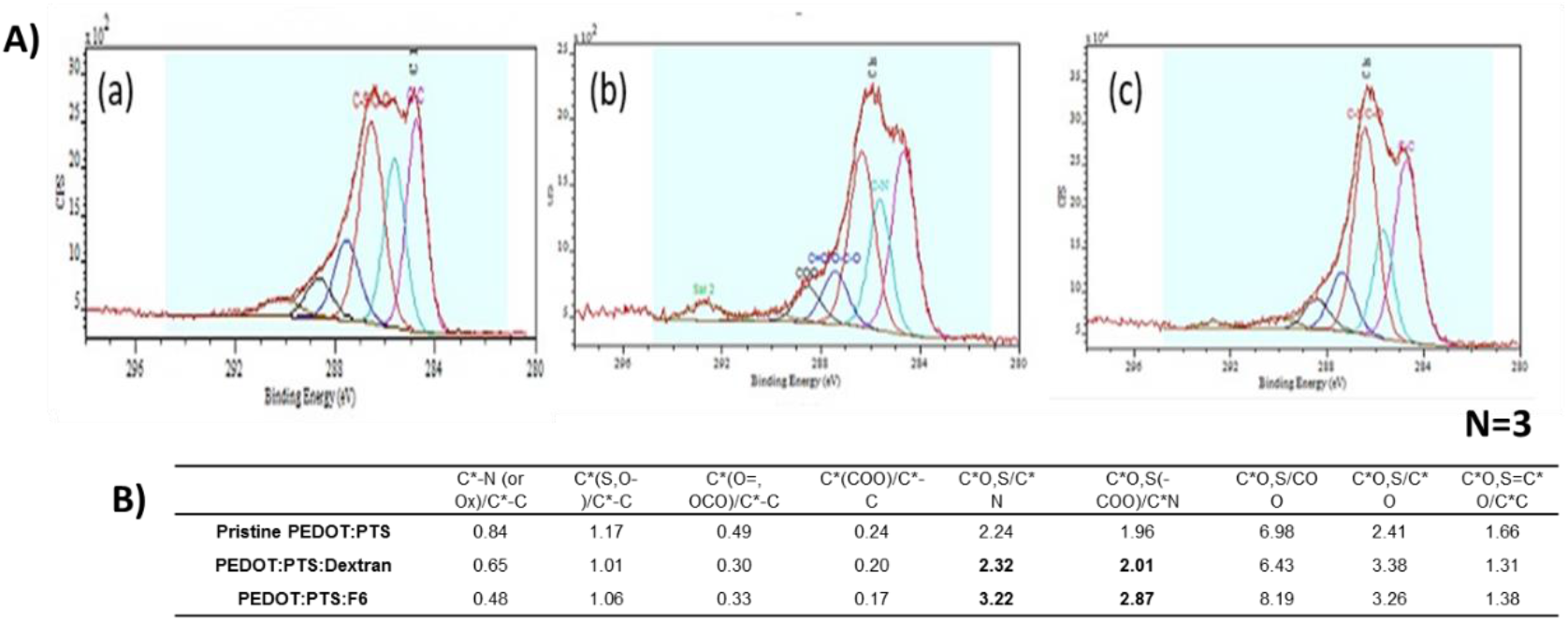
The positions of C peaks and the relative compositions. A) X-ray photoelectron spectra of (a) pristine PEDOT:PTS coated microelectrode, (b) PEDOT:PTS:Dextran coated microelectrode, and (c) PEDOT:PTS:F6 coated microelectrode showing the presence of carbon in various oxidation states. The C-N bond cannot be used to discern the percentage of alanine moiety present as a substitution on heparan sulphate mimetic-F6, because trace amounts of acetonitrile from the electrolyte could be present contributing to this peak in all three coated microelectrodes. C* denotes partially oxidised carbon. Table giving the relative atomic percentages of these bonds in coated microelectrodes is presented in B) the relative increase in the percentages of C*O,S/C*N and C*O,S(-COO)/C*N bonds between PEDOT:PTS:Dextran and PEDOT:PTS:F6 (highlighted in bold), shows the increase in N possibly due to presence of phenylalanine component in F6.

The data presented above reports on the super-hydrophilic chemical character of the PEDOT:PTS:F6 coated microelectrodes owing to the successful incorporation of F6 compared to pristine PEDOT:PTS coatings. This favorable chemical surface character of the PEDOT:PTS:F6 coating is important for material performance towards the development of mimetic neural coatings.

#### 2.2.2 Electrochemical Characterisation

The selection of dopants and their ionic nature are of particular importance in electrochemical deposition, and the formulation of electrically stable conducting polymers as neuroelectrode coatings [33, 90–94] and ultimately impact upon the materials physico-chemical, electrical and biological characteristics [33, 95].

Although a number of studies have shown that the incorporation of biomolecules as dopants within a conducting polymer can initiate biological response in vitro [2, 25, 27, 28, 34, 36, 96], a majority of these studies also note an associated deterioration in the conducting polymer electrochemical properties relative to pristine polymers or polymers doped with inorganic ionic species [32, 94, 97, 98]. Research to date on the electrochemical effects of doping conducting polymers with biochemical molecules has not been extended to the effect of HS mimetic incorporation into PEDOT derivatives. However, initial physicochemical analysis indicated that PEDOT:PTS:F6 electrode coatings presented a significantly rougher, super-hydrophilic surface relative to pristine PEDOT:PTS coating, suggesting that Pt/Ir microelectrodes coated with this chemistry would exhibit an enhanced charge transfer capability and neural stimulation efficacy.

To assess the electroactivity and electrochemical stability of electrodeposited chemistries, CV was performed with PtIr, pristine PEDOT:PTS and PEDOT:PTS:F6 coated microelectrodes, as recorded in PBS with microelectrodes possessing a surface area of 0.054 mm^2^ (***Figure 11*A**). CV curves of PEDOT:PTS:F6 coated microelectrodes demonstrated defined oxidation – reduction peaks and demonstrated a strong capacitive behavior, especially within the potential range from −0.2 V to 0.8 V (vs. Ag/AgCl), when compared to PEDOT formed in the presence of conventional sulphonate anion PTS only.

**Figure 11.**
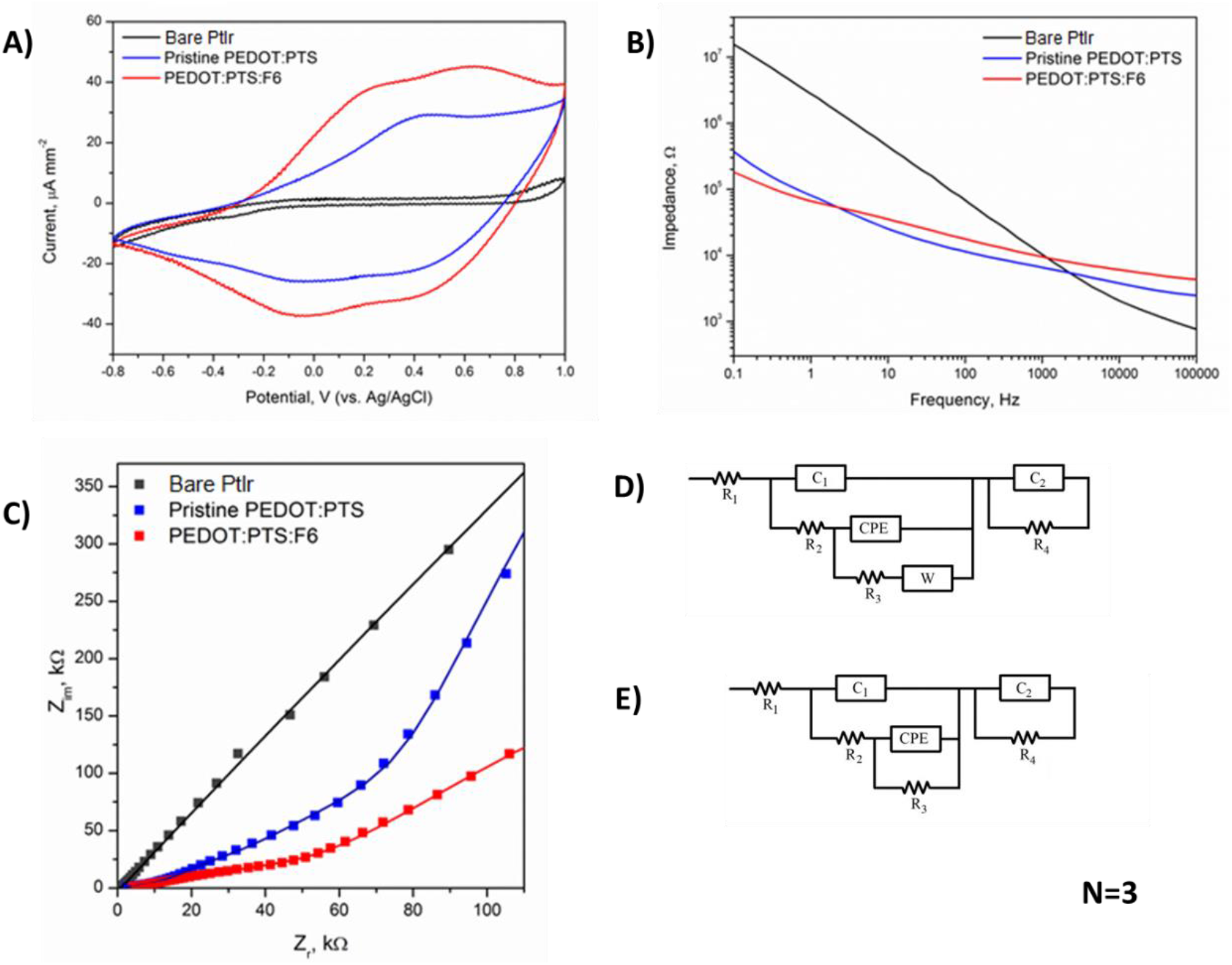
Electrochemical analysis of PEDOT:PTS:F6 coated microelectrodes. A) Cyclic voltammograms (CVs) of bare Platinum/Iridium (PtIr) microelectrodes, pristine PEDOT:PTS and PEDOT:PTS:F6 coated microelectrodes. CVs were recorded in 0.01 M phosphate-buffered saline (PBS) at a scan rate of 100 mV s^−1^. Bode B) and Nyquist C) plots comparing the EIS spectra of bare Platinum/Iridium (PtIr) microelectrodes, pristine PEDOT:PTS, and PEDOT:PTS:F6 coated microelectrodes. D) and E) represent the electrical equivalent circuit used to analyse experimental data of the coated microelectrodes and bare Platinum/Iridium (PtIr) microelectrodes, respectively. Results are ± STD.

As reported previously [99–101], PEDOT coating have a strong beneficial effect on the CSC of metallic microelectrodes, and a significant increase in the CSC was observed following electrodeposition of PEDOT:PTS onto Pt/Ir microelectrodes, from 99.7 ± 17.2 μC mm^−2^ for bare Pt/Ir microelectrodes to 697.9 ± 23.7 μC mm^−2^ for pristine PEDOT:PTS coated microelectrodes (***Table 7***). The incorporation of F6 induced a further significant increase in the CSC of coated Pt/Ir microelectrodes, to a value of 899.5 ± 1.6 μC mm^−2^. Critically this observed CSC was substantially higher than those recorded with other conducting polymer-based systems employed in neural applications, e.g. surfactant-templated ordered PEDOT (26.7 mC cm^−2^) [102], PEDOT-CNT (70 mC cm^−2^) [103] and PEDOT:PSS (75.6 mC cm^−2^) [100].

**Table 7.** Electrochemical performance of PEDOT:PTS:F6 coated microelectrodes. The charge storage capacity (CSC) was evaluated from the cathodic region of cyclic voltammograms (CVs) recorded in 0.01 M phosphate-buffered saline (PBS) at 100 mV s^−1^ scan rate (potential range: −0.8 V to 1 V vs. Ag/AgCl). The calculated resistance values (R3) of bare Platinum/Iridium (PtIr) microelectrodes, pristine PEDOT:PTS and PEDOT:PTS:F6 coated microelectrodes after equivalent circuit analysis. Results are ± STD. N=3

| | CSC,<br>[$\mu\text{C mm}^{-2}$] | R<sub>3</sub><br>[kΩ] |
| --- | --- | --- |
| **PtIr** | 99.7 ± 17.2 | 591.9 ± 144.8 |
| **Pristine PEDOT:PTS** | 697.9 ± 23.7 | 41.9 ± 2.1 |
| **PEDOT:PTS:F6** | 899.5 ± 1.6 | 17.0 ± 3.5 |

Impedance spectra of PEDOT:PTS:F6 and pristine PEDOT:PTS coated microelectrodes and a control Pt/Ir microelectrode group were obtained via a three electrode set-up to assess the effects of F6 doping on film resistance ***Figure 11*B**. The Bode plots indicates PEDOT:PTS:F6 coated microelectrodes exhibited the lowest impedance profile in the low frequency range (below 1 Hz). For the rest of the evaluated frequencies (1 Hz to 100K Hz) PEDOT:PTS:F6 coated microelectrodes and pristine PEDOT:PTS coated microelectrodes show similar profiles, and a significant decrease in impedance relative to uncoated microelectrodes in the 0.1 Hz to 1K Hz range. However, due to the complex character of the impedance response, which is typical for microelectrodes and microelectrode arrays [104], Nyquist plots were generated to compare the resistance of the electrodes (***Figure 11*C**). These plots display both amplitude and phase angle on a single plot using frequency as a parameter, and, when fitted with an appropriate equivalent electrical circuit, can be used to provide the quantification of electrical properties of all elements of the circuit. Through detailed analysis of the Nyquist plots, it was possible to report on the charge transport mechanism and to extract solution resistance from the overall impedance profile and study only the parameters related to the properties of electrodeposited coatings [36]. The equivalent circuit for pristine PEDOT:PTS and PEDOT:PTS:F6 coated microelectrodes included bulk solution resistance of the polymer and the electrolyte (R_1_), double layer capacitance (C_1_), resistance of the electrolyte (R_2_), constant phase element (CPE), charge transfer (R_3_) and Warburg impedance of the polymer (W), second capacitor element (C_2_) as well as charge transfer resistor (R_4_), (***Figure 11*D**) as described previously [105]. In the case of bare PtIr microelectrodes, the equivalent circuit was the same except for the Warburg impedance element (***Figure 11*E**). The simulation data confirmed the lowest resistance profile of PEDOT:PTS:F6 coated microelectrodes (17.0 ± 3.5 kΩ), when compared to pristine PEDOT:PTS (41.9 ± 2.1 kΩ) and bare PtIr (591.9 ± 144.8 kΩ) microelectrodes (***Table 7***).

It can be hypothesised that F6 acted as a stable dopant for PEDOT, as observed with electrodeposited PEDOT:PSS, for which positively charged PEDOT oligomers are attached to the negatively charged, high molecular weight PSS, resulting in the stacked arrangement of polythiophene rings responsible for high conductivity of the material [106, 107]. Similarly, it can be hypothesised that the large F6 counterion enhances the self-arrangement of growing PEDOT chains with more defined oligomers than occurs in the presence of PTS alone [15, 106, 107]. It can also be argued that the morphological and super-hydrophilicity properties of the PEDOT:PTS:F6 coated microelectrodes, further increase the availability of sites for charge transfer and contribute to the observed improvement in CSC of PEDOT:PTS:F6 coated microelectrodes relative to pristine PEDOT:PTS coated microelectrodes which are represented by the increased conductivity of the PEDOT:PTS:F6 coated microelectrodes [15, 81, 108]. The benefit of F6 as the counterion for the functionalised PEDOT:PTS:F6 coating is here shown with the potential of a structure-electronic surface that lays foundation for future coating-electrode design.

#### 2.2.3 Biological Characterisation

The aim of reducing inflammation while promoting neural tissue integration and maintaining device functionality represents a paradgime of the field of bio-interface engineering [21, 30, 97, 109–115]. GAGs, as key components of the ECM architecture, and as charged biomolecules that can act as counterions in the electrochemical polymerisation of conducting polymers, represent ideal dopants for the formulation of functional conducting polymer interfaces to modulate cellular function and tissue homeostasis [116, 117].

In particular, the HM F6 has been shown previously to present efficacy in the regulation of neurodegenerative diseases including the tauopathies and synucleinopathies *in vitro* and *in vivo* by blocking the internationalisation and propagation of protein aggregates [60]. Therefore, it was the hypothesis of this work that through the incorporation of the F6 as co-dopant for EDOT electrical polymerisation biomimetic coatings could be developed to promote neural integration and minimise the inflammatory response associated with reactive gliosis *in vitro*.

Initially, to determine a release profile of the HS mimetic F6 from the PEDOT:PTS polymer and to understand the bioactivity of the F6 in subsequent *in vitro* culture studies, the elution profile of PEDOT:PTS:F6 coated electrodes was assessed through a spontaneous passive mode over a period of 21 days simulating *in vivo* conditions (constant aggitation under 37° degrees), (***Figure 12***). The spontaneous elution profile shows an initial burst of the F6 taking place within the first three days of the experiment with observed eluted F6 concentrations of 5.9 µg ml^−1^ cm^−2^, followed by a slow increase by day seven to reach a plateau with a released concentration of F6 of 11.1 µg ml^−1^ cm^−2^ by day twelve. This behavior is in agreement with the commonly observed pattern of spontaneous passive release of biomolecules from polymeric matrices observed by others [21, 118, 119], a trend associated with the elution of the active molecules from the near-surface region of the matrix.

**Figure 12.**
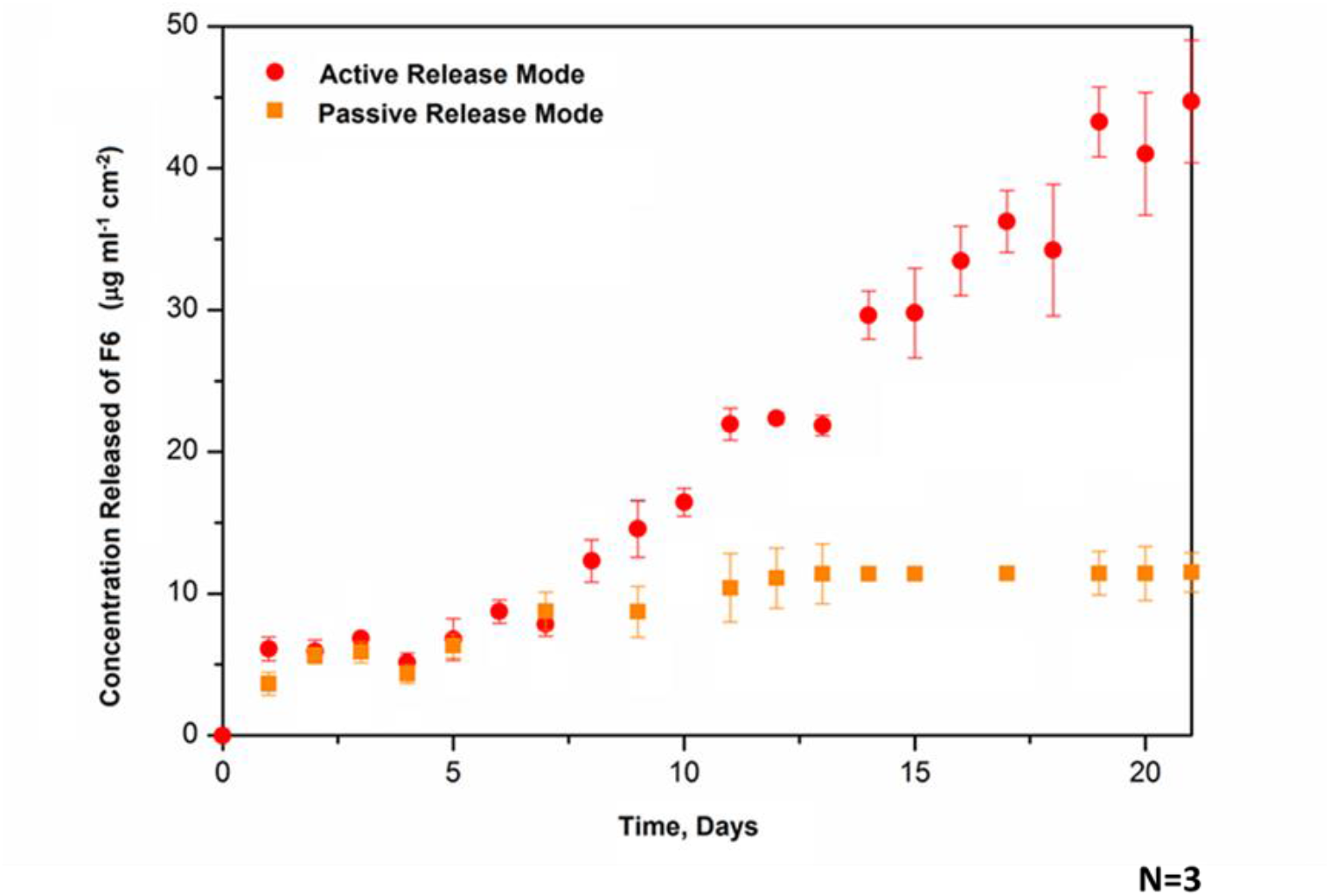
Concentration profile of heparan sulphate mimetic-F6 released from PEDOT:PTS:F6 coated electrodes in the passive (orange squares) and active (red dots) modes for a period of 21 days under simulating in vivo conditions (constant aggitation under 37° degrees). For the active release mode, 50 stimulation cycles were applied every day over 21 days which resulted in approx. 1000 stimulations over this period. The stimulation regime was a biphasic potential pulse, consisting of a 5-second application of a reduction potential (−0.5 V) followed by a 5-second application of an oxidative potential (+0.5 V). Results are ± STD.

Further, and owing to the excellent electrochemical properties of the PEDOT:PTS:F6 coatings, these results were compared to an electrically active mode of F6 release from bio-functionalised coated electrodes to evaluate the effective release of F6 in response to a therapeutically relevant voltage pulse (***Figure 12***). PEDOT:PTS:F6 coated microelectrodes were subjected to a biphasic potential pulse of 5-second duration to apply a reduction potential (−0.5 V) followed by a 5-second application of an oxidative potential (+0.5 V). Stimulation conditions mimicked those of the passive study but under a stimulation regime of 50 cycles (corresponding to 500 s) every day over a period 21 days (approx. 1000 stimulations in total) [31, 120–122].

In contrast to the spontaneous passive release of F6, with electrically triggered PEDOT:PTS:F6 coated electrodes the concentration of F6 increased linearly over time and to reach a concentration of 44.7 µg ml^−1^ cm^−2^ by day 20, a concentration that was shown to promote high cell viability (***Figure 4***) when added unbound into culture. Interestingly, the F6 released from electrically active electrodes exhibited a release profile similar to that of the electrically passive electrodes until day seven. Following this initial period bi-phasic stimulation induced a linear release of significantly higher amounts of F6 relative to electrically passive electrodes. It follows, that by modification of the electrode stimulation parameters, it could be possible to temporally control the release profile of F6, to maximise the therapeutic potential *in vivo* or to prolong the elution period [120], important considerations for future work into developing functionalisation approaches for long term applications.

To further assess stability of the electrochemical performance of the coated microelectrodes, pristine PEDOT:PTS and PEDOT:PTS:F6 coated microelectrodes and bare Pt/Ir microelectrodes were subjected to 1000 continuous stimulation cycles corresponding to the cumulative stimulus used for the active release mode of the F6. The final performance of the microelectrodes was evaluated in terms of changes in charge storage capacity after stimulation as detailed in ***Figure 13***. Electrodes exhibited maintained electrical integrity after electrical stimulation conditions, with the incorporation of the F6 within the PEDOT matrix.

**Figure 13.**
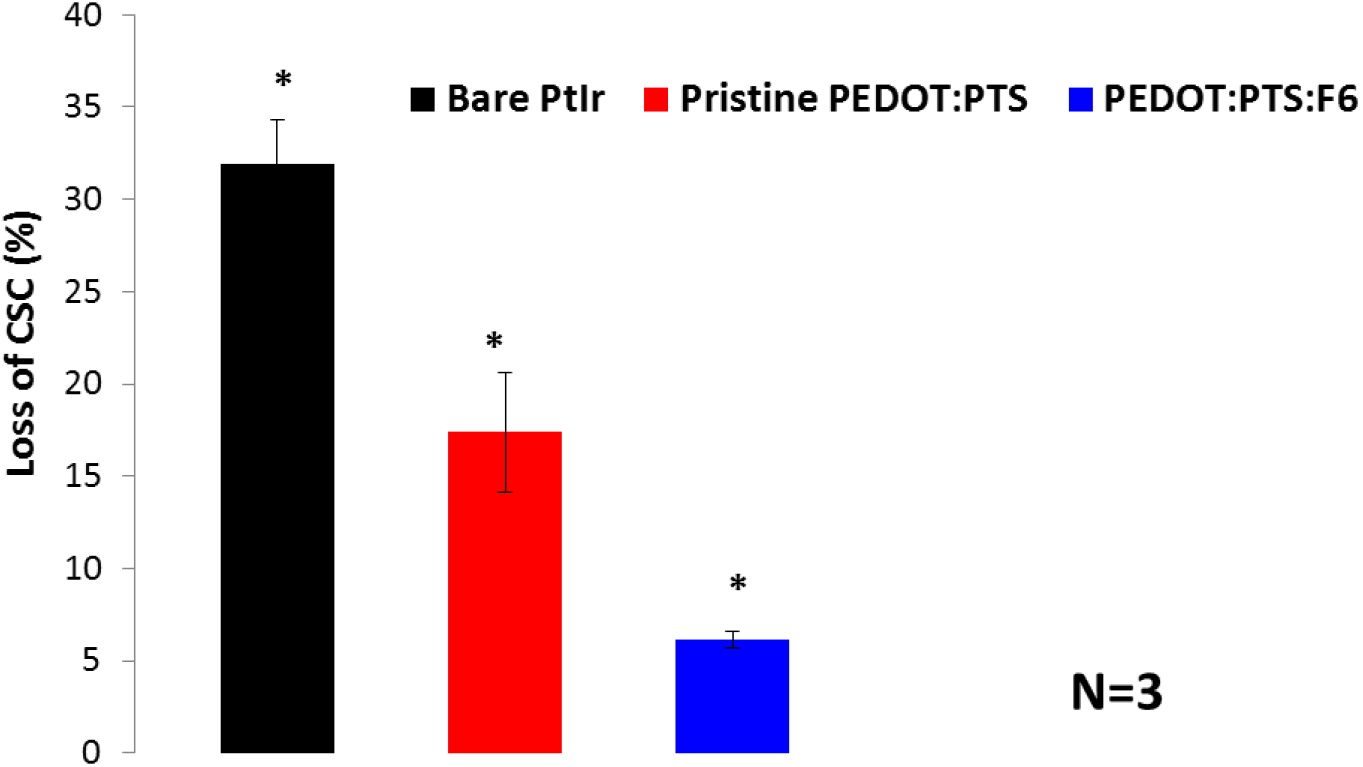
Electrical stability of the PEDOT:PTS:F6 microelectrodes. Plot of percentage loss of charge storage capacity (CSC) after 1000 continuous stimulation cycles of biphasic potential pulses, consisting of a 5-second application of a reduction potential (−0.5 V) followed by a 5-second application of an oxidative potential (+0.5 V). Important significant differences between the bare PtIr microelectrodes and the PEDOT-based coated microelectrodes were observed. PEDOT:PTS:F6 coated microelectrodes were the most stable followed by pristine PEDOT:PTS coated microelectrodes and bare PtIr microelectrodes after. Results are ± STD, ★ = p< 0.05.

The bioactivity of PEDOT:PTS:F6 coatings was evaluated *in vitro* with planar 1.6 cm^2^ platinum (Pt) coated electrodes using a primary ventral mesencephalic (VM) mixed neural cell population. A non-sulphated polymeric dextran was also used as negative control to understand the effect of the sulphation pattern provided by the mimetic on cell function. A sputtered platinum (Pt) group was also used in the in vitro studies as a representative electrode control material.

***Figure 14*A** shows representative fluorescent micrographs of VM derived mixed cultures of neurons and astrocytes cultured on experimental and control electrodes. All groups were evaluated over a period of three, seven and ten days in culture. The persistence of neurons and astrocytes on each of the experimental and control groups as a function of time is presented in ***Figure 14*B**. It was evident that electrodes subjected to the electrodeposition of PEDOT:PTS doped with non-sulphated polymeric dextran (PEDOT:PTS:Dextran) induced an overall marked increase in astrocyte density, accompanied with a significant decrease in neuronal presence as a function of time. A trend that contrasted significantly to all other experimental groups and to Pt control substrates. By day ten, the PEDOT:PTS:Dextran coated electrodes exhibited an astrocytes presence of 92.86 % and a neurons presence of 7.15 %. In contrast, a maintained significant lower presence of astrocytes on PEDOT:PTS:F6 coated electrodes over that of Pt control and pristine PEDOT:PTS coated electrodes was consistently observed over time, findings that were coupled with an interesting and significantly higher presence of neural density on these electrodes at each time point relative to all the experimental and control groups. By day ten, the recorded percentage of astrocytic cell density on the PEDOT:PTS:F6 coated electrodes was 42.86 % with a percentage of neuron presence of 57.14%, followed by pristine PEDOT:PTS coated electrodes with a 46.39 % astrocytes presence and a 53.60 % neuron presence and Pt controls with 61.97% astrocytes presence and 38.03% neural density.

**Figure 14.**
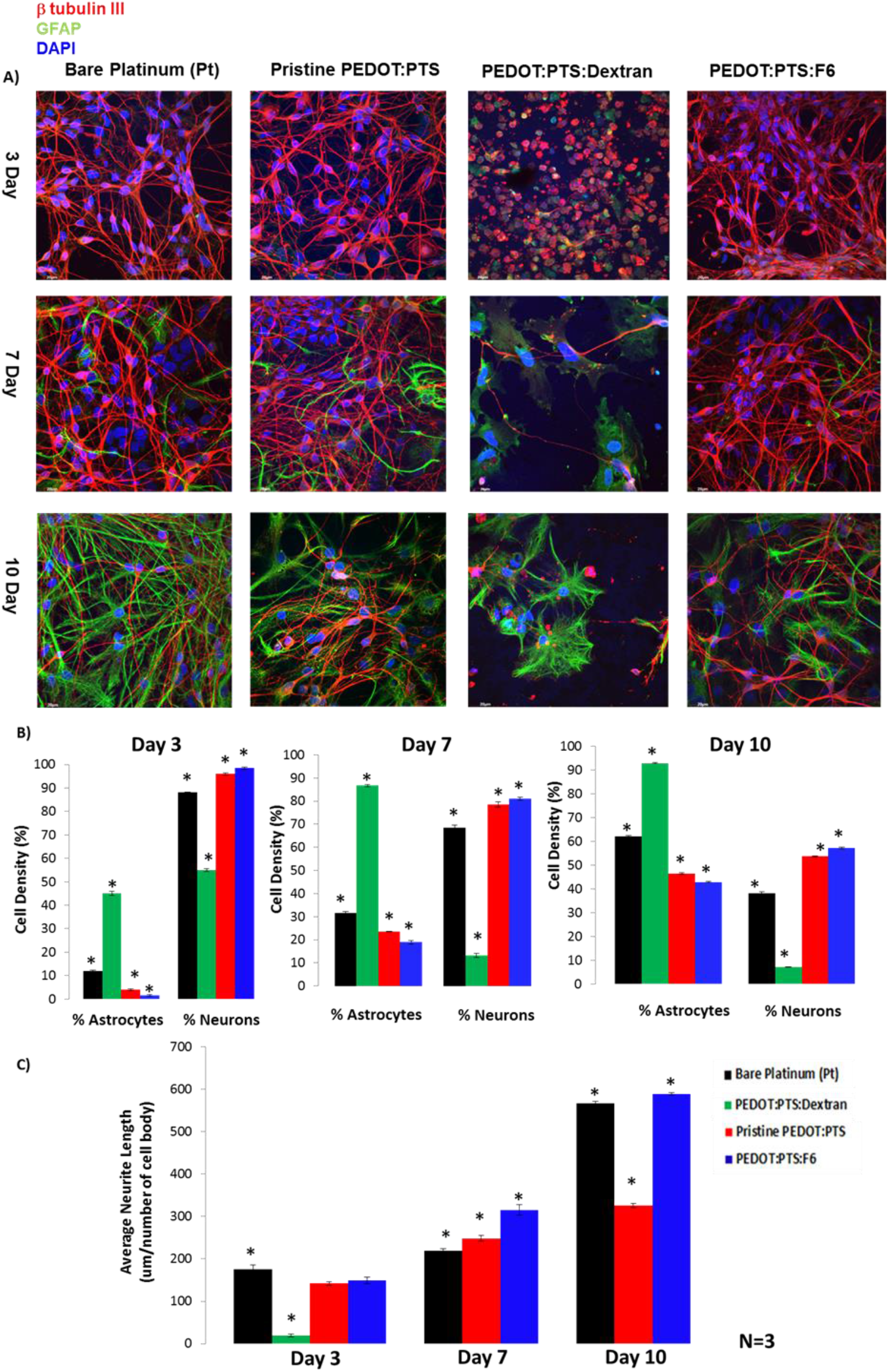
Cytocompatibility of PEDO:PTS:F6 coated electrodes. A) Fluorescent images of primary ventral mesencephalic (VM) mixed cell population grown on each of the bare platinum (Pt) electrodes and pristine PEDOT:PTS, PEDOT:PTS:Dextran and PEDOT:PTS:F6 coated electrodes for three, seven and ten days in culture. Neurons are visualised by anti β-tubulin III, in red, astrocyte cells by anti-GFAP, in green and nuclei are visualised by DAPI, in blue. Bar = 20 µm, objective 60× magnification. Cell density (%) analysis of astrocytes and neurons presence on each of the electrodes is presented in B). An overall significant (p < 0.05) enhancement in viability of neurons was observed in PEDOT:PTS:F6 coated electrodes. Neural length analysis of electrodes presented in C) showed with significant (p < 0.05) longer neurite lengths the neurons grown on PEDOT:PTS:F6 coated electrodes, suggesting on an overall neurotrophic effect of these electrodes owing to the presence of heparan sulphate mimetic-F6. Results are ± STD. ★ = p< 0.05.

In conjunction with cell density, as an indicator of cell viability, neurite length on experimental and control substrates was also analysed (***Figure 14*C**). At day three, similar neural lengths were observed in neurons cultured on pristine PEDOT:PTS coated electrodes and PEDOT:PTS:F6 coated electrodes with neural lengths of 142.21 µm ± 4.21 and 149.60 µm ± 7.45 respectively. Neurite length was significantly lower than those of neurons cultured on Pt electrodes on which mean neurite length was of 175.08 µm ± 10.55, but significantly higher than the neural length observed on the PEDOT:PTS:Dextran coated electrodes of 18.85 µm ± 3.15. This trend was lost by day seven and neurite lengths were significantly increased on PEDOT:PTS:F6 coated electrodes relative to Pt control and PEDOT:PTS coated electrodes. However, neurite length on the non-sulphated polymeric dextran PEDOT:PTS:Dextran control samples could not be quantified at the subsequent time points due to a significantly reduced neuron presence, rendering the application of the stereology method used for the quantification of length invalid.

Additional significant differences in neural length were observed by day ten, with significant neurite elongation exhibited in cells cultured on the PEDOT:PTS:F6 coated electrodes (589.20 µm ± 2.90) relative to bare platinum control (566.36 µm ± 6.35) and pristine PEDOT:PTS coated electrodes (325.76 µm ± 4.66).

These results show that the PEDOT:PTS:Dextran coated electrodes induced overall poor viability of the primary VM neural population relative to pristine PEDOT:PTS coated electrodes and functionalised PEDOT:PTS:F6 coated electrodes, resulting, as a consequence, in an over proliferation of astrocytes. Moreover, this finding has implications for the role of sulphation on the modulation of neural survival, as seen by others [36, 123–125]. In fact the incorporation of the F6 within the PEDOT:PTS polymer matrix showed that these coatings do not enhance astrocyte proliferation and adhesion relative to control Pt and to pristine PEDOT:PTS coated electrodes for up to ten days in culture, rather, the presence of F6 significantly enhanced neuron survival and supported neurite extension of the mesencephalic neuros *in vitro* relative to all experimental and control groups. Therefore, a neurotrophic effect imparted by F6 incorporated in the PEDOT:PTS polymer matrix is suggested.

Further, the synthesis of pro-inflammatory cytokines from VM populations was assessed *in vitro* to support the inflammatory modulatory effect of F6 when incorporated in the PEDOT:PTS matrix. Chemokines IFN-g, TNF-α, IL-6, IL-5, and KC/GRO were selected for analysis, signaling factors involved in mediating neuronal-glial interactions [126, 127] [65, 128–132]. The release profiles were compared across the experimental groups, controls and with an additional inflammatory control group, VM cells cultured on Thermanox® Plastic Coverslips (tissue culture plastic) which received a stimulus of interleukin-1beta (IL-1β) at a dose of 10 ng ml^−1^.

The secretion of pro-inflammatory cytokines and chemokine factors in mixed cell populations cultured on all experimental and control groups is presented in ***Figure 15*A-E**.

**Figure 15.**
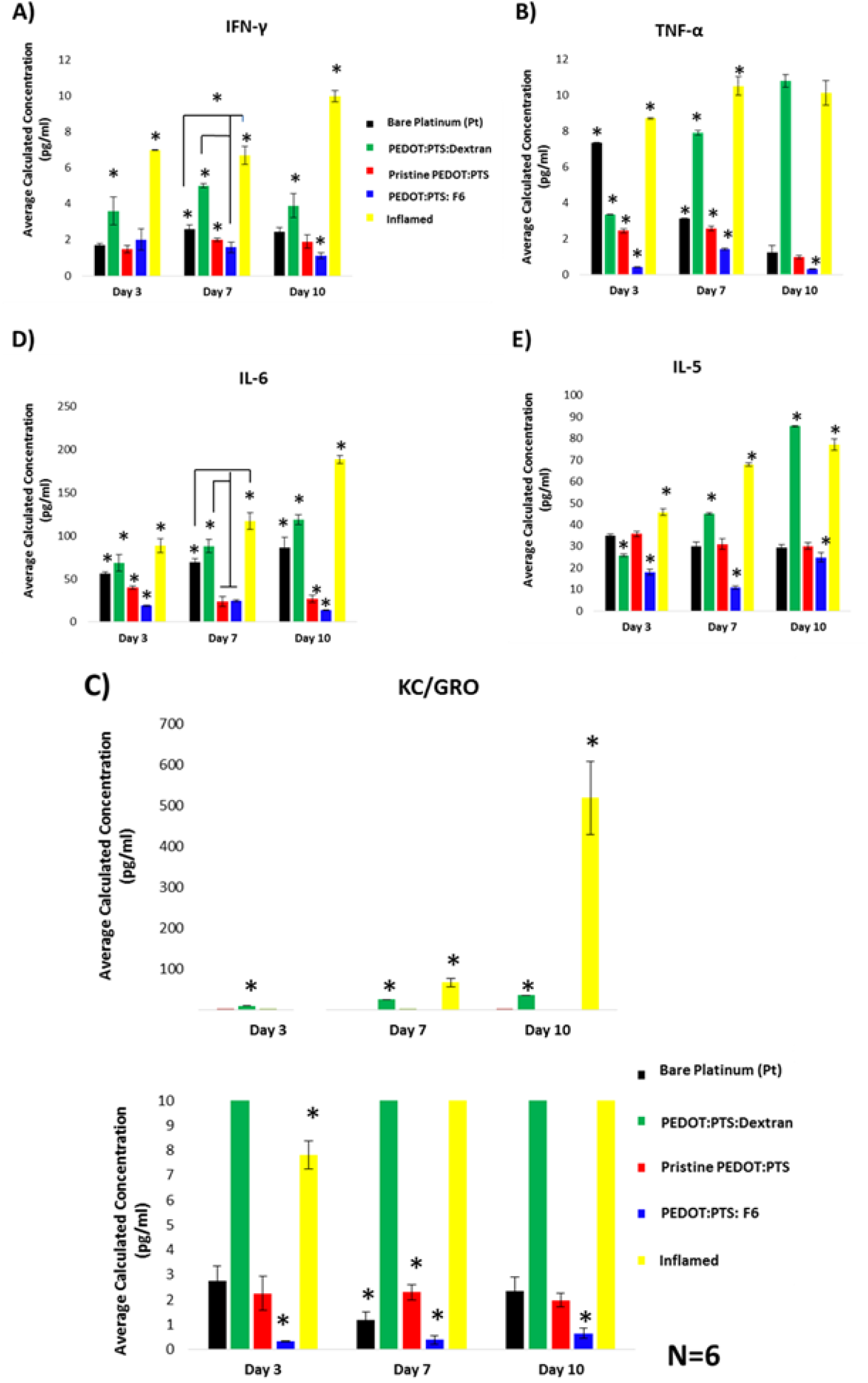
Comparative pro-inflammatory derived cytokines and chemokine factor profiling for interferon-γ (IFN-γ;A), tumor necrosis factor-α (TNF-α;B), chemokine factor CXCL-1 (KC/GRO;C), interleukin-6 (IL-6;D) and interleukin-5 (IL-5;E). The release expression of each of the signaling molecules is analysed from primary ventral mesencephalic (VM) mixed cell population supernatants collected at day three, seven and ten days on bare platinum (Pt) electrodes, pristine PEDOT:PTS, PEDOT:PTS:Dextran and PEDOT:PTS:F6 coated electrodes. An important effect is seen on the PEDOT:PTS functionalised with the heparan sulphate mimetic-F6 coated electrodes, which presented significantly low release profiles for each cytokine and chemokine factor analysed after ten days in culture. Results are ± STD, ★ = p< 0.05.

***Figure 15*A-B** show the release profiles of IFN-g and TNF-α, respectively. These cytokines are known to act in synergy [133–135], and showed a similar release profile. It was observed that the release of IFN-g from VM cells cultured on PEDOT:PTS:F6 coated electrodes underwent a linear decrease with time, and was significantly lower by day ten relative to IFN-g synthesis on Pt control, PEDOT:PTS based-coated electrodes, and inflamed control, respectively. Interestingly, the IFN-g release profile of VM cells cultured on Pt and pristine PEDOT:PTS coated electrodes was maintained at comparatively similar levels by day ten. Further, PEDOT:PTS:Dextran coated electrodes induced the second highest VM release profile of IFN-g after that of cells cultured on the inflamed control. Conversely, TNF-α release profiles showed an overall linear decrease with time from VM cells cultured on bare Pt electrodes, pristine PEDOT:PTS and PEDOT:PTS:F6 coated electrodes, yet cells cultured under inflamed conditions and on PEDOT:PTS:Dextran control groups presented a consistent increase in TNF-α release. By day ten, a significantly higher release profile of TNF-α was observed with VM cells cultured on the PEDOT:PTS:Dextran coated electrodes which was comparable to that observed in VM cells cultured under inflamed control conditions. Control Pt, pristine PEDOT:PTS and PEDOT:PTS:F6 groups, were associated with a significant reduction in TNF-α synthesis, with an overall significantly lower TNF-α expression by the cells cultured on the functionalised PEDOT:PTS:F6 electrodes.

VM cell expression of the chemoattractive factor CXCL1 [136, 137] also known as KC/GRO, was significantly reduced at all time points when cultured on PEDOT:PTS:F6 coated electrodes relative to all experimental and control groups (***Figure 15*C**). In contrast, a significantly higher production of KC/GRO was observed with control PEDOT:PTS:Dextran coated electrodes and with the inflamed group.

Further, an increase in the secretion of cytokines IL-6 [138, 139] (***Figure 15*D**) and IL-5 [140–142] (***Figure 15*E**), as result of the co-stimulatory cross-talk of cell-cell interactions produced by the nature of the mixed VM populations was observed, indicating the biological protection offered by the PEDOT:PTS:F6 coated electrodes relative to control Pt electrodes, pristine PEDOT:PTS and PEDOT:PTS:Dextran coated electrodes, and inflamed control respectively. Also, the release expression of IL-6 and IL-5 by the VM populations cultured on the PEDOT:PTS:Dextran coated electrodes, reflects on the overall promotion of a more pro-inflammatory milieu, which relates to the poor VM cell viability and neural outgrowth observed on these electrodes.

The modulation of cytokine and chemokine activity in VM populations cultured on PEDOT:PTS functionalised with F6 indicates the protective effect of F6 on matrix proteins from protease activities [35, 44], which translates to an enhanced availability [40, 54, 65, 137] of neurotropic growth factors.

Considerable research has shown that the ECM presents a spatial network of functional GAGs moieties located on proteoglycans which are present at the cell surface [143–145] and exert biological activities by interacting with a broad spectrum of protein ligands including growth factors, cytokines and ECM proteins [145]. The molecular mechanism of GAGs like HS that mediated cell signaling has been extensively studied, and a high binding affinity for FGF-2 promotes the formation of a stable tertiary signal complex [146]. HS-protein binding processes are known to be mediated by specific sulphation patterns that recognise protein binding sites [147] and promote cellular signaling processes including cell proliferation and differentiation [73] and more specifically, they play essential roles in inflammation [65, 148], wound healing [149], angiogenesis [150] and anticoagulation [151, 152]. To note, all these processes are not mediated by a single generic HS polysaccharide, but by many different HS species which differ in sulphated disaccharide profile. Here F6, a chemically modified molecule from the HM library was synthesised with a 0.7 (70%) statically available sulphate substitution (dsS) along the saccharidic chains of a sulphation pattern on 2-0-sulphated in percentage (***Figure 3***). These structural properties of F6 have particular important correlations with the findings within this work and it can be hypothesised that the neurotrophic and neuroprotective effects observed with PEDOT:PTS:F6 coated electrodes were potentiated through binding with the anionic characteristics of F6. In consequence, potentialise the binding with secreted neurotropic growth factors such as FGF-2 [144, 145, 153] and VEGF [154, 155].

FGF-2 has been shown to promote neural survival and the development of ventral midbrain neurons with a neurotrophic function that enhances neural networking, neural outgrowth and guidance [156–159]. Moreover, VEGF has been shown to promote neuroprotection indirectly by activating the proliferation of glia and by promoting neurogenesis and neuronal patterning in mixed embryonic VM cultures [160, 161].

Therefore, as a proof of concept study, the binding capacity of F6 with FGF-2 and VEGF were assessed with a competitive binding affinity assay as previously described [73] (***Figure 16***). The binding potential of F6 and other exogenous GAGs HS and heparin to FGF-2 and VEGF was assessed within the limits of F6 concentrations that were previously shown to maintain high cell viability (***Figure 4***) when added unbound into the culture medium. Heparin was used as a biochemical control (100% relative effect), and naturally occurring HS as relevant physiological control. Dextran was further used as a non-sulphated GAG control.

**Figure 16.**
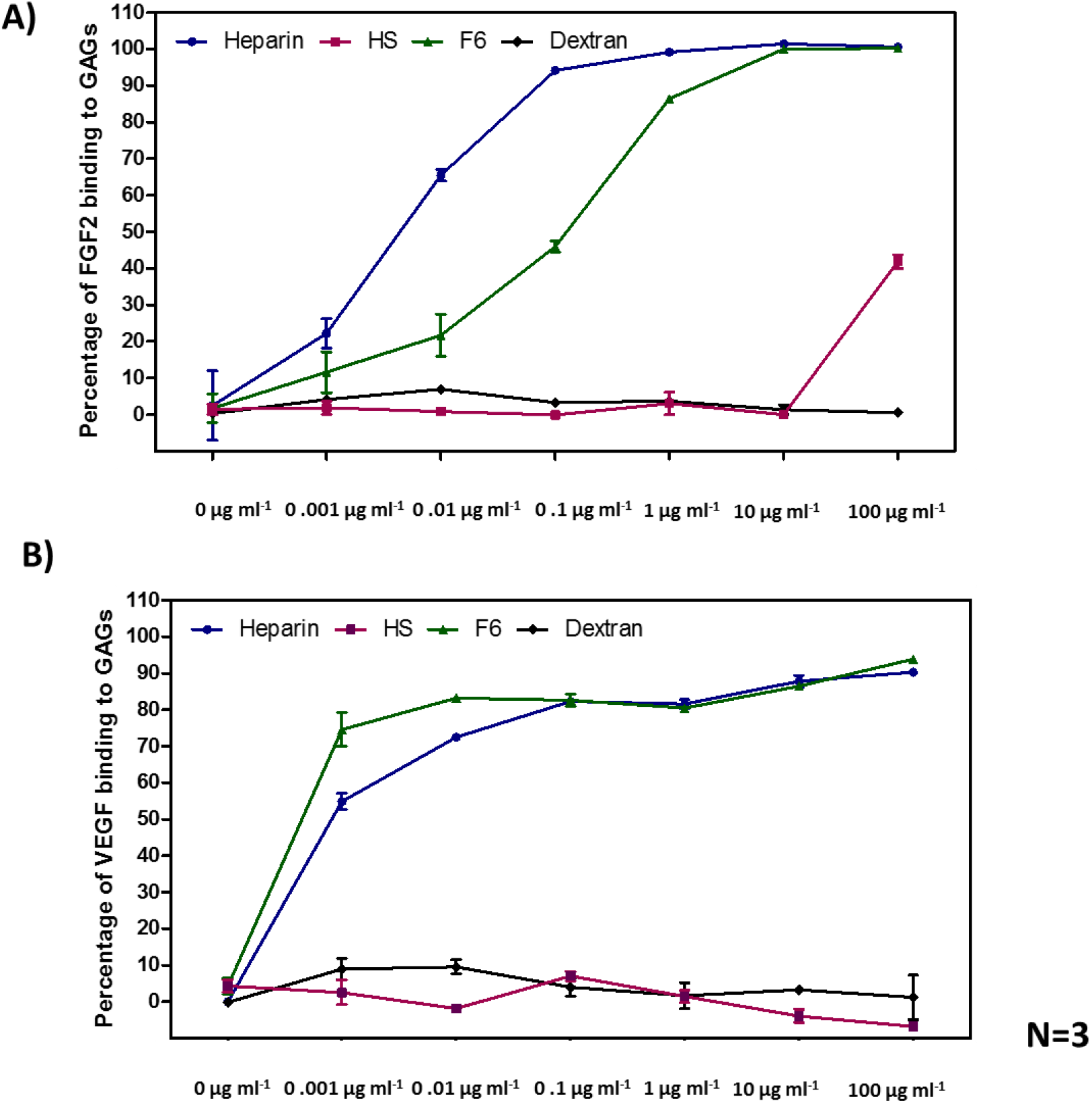
Glycosaminoglycans competition assay towards FGF-2 and VEGF binding. F6 shows a strong binding capacity to FGF2 and VEGF in a heparin-like manner.

***Figure 16* (A-B)** shows the binding of F6 to FGF-2 and VEGF respectively. It is interesting to observe that F6 binds to both growth factors in a heparin-like manner, suggesting the presence of structural binding motifs with 2-O-sulfated and 6-O-sulfated domains for the binding with FGF-2 and VEGF growth factors, respectively [162–165]. But further, this data suggests that F6 adopts a favourable molecular conformation in solution that facilitate the recognition of important biological growth factors compared to the physiological control HS. The introduction of hydrophobic moieties in polyanions, as is the case of the L-phenylalanine methyl ester in F6, can be expected to increase their affinity for proteins and participate in the control of the conformational changes associated to pathogenesis [78]. Our results suggest that there is a structural rational for the binding potential of F6 to the studied growth factors. This rational can be used to analyse these activities through recognition studies using molecular and conformational analysis.

Overall, these findings are particularly important for neuroelectrode coating designs with biological dopants as they not only show an overall reduction of inflammation profiling while promoting neural growth through a mechanistic binding with growth factors *in vitro*, but further open up the study of glycan mimetics to potentiate matrix-therapeutics strategies at the neural-tissue interface.

Complementary to the cytokines-protecting response, and as part of a synergistic approach with an in-house gliosis antibody microarray as previously detailed in [5], the neuroprotection effect of the PEDOT:PTS:F6 coated electrodes was further evaluated in the context of gliosis, a process shown to decrease adjacent neurons from the electrode *in vitro* and *in vivo* conditions [5–9]. Details of the antibodies used and their corresponding optimised concentrations are in ***Table 1***.

Unsupervised clustering of the expression of pro-gliosis proteins was performed with VM cells cultured on all experimental and control groups and an additional inflamed control for three days (***Figure 17*A**) and for ten days *in vitro*. (***Figure 17*B**).

**Figure 17.**
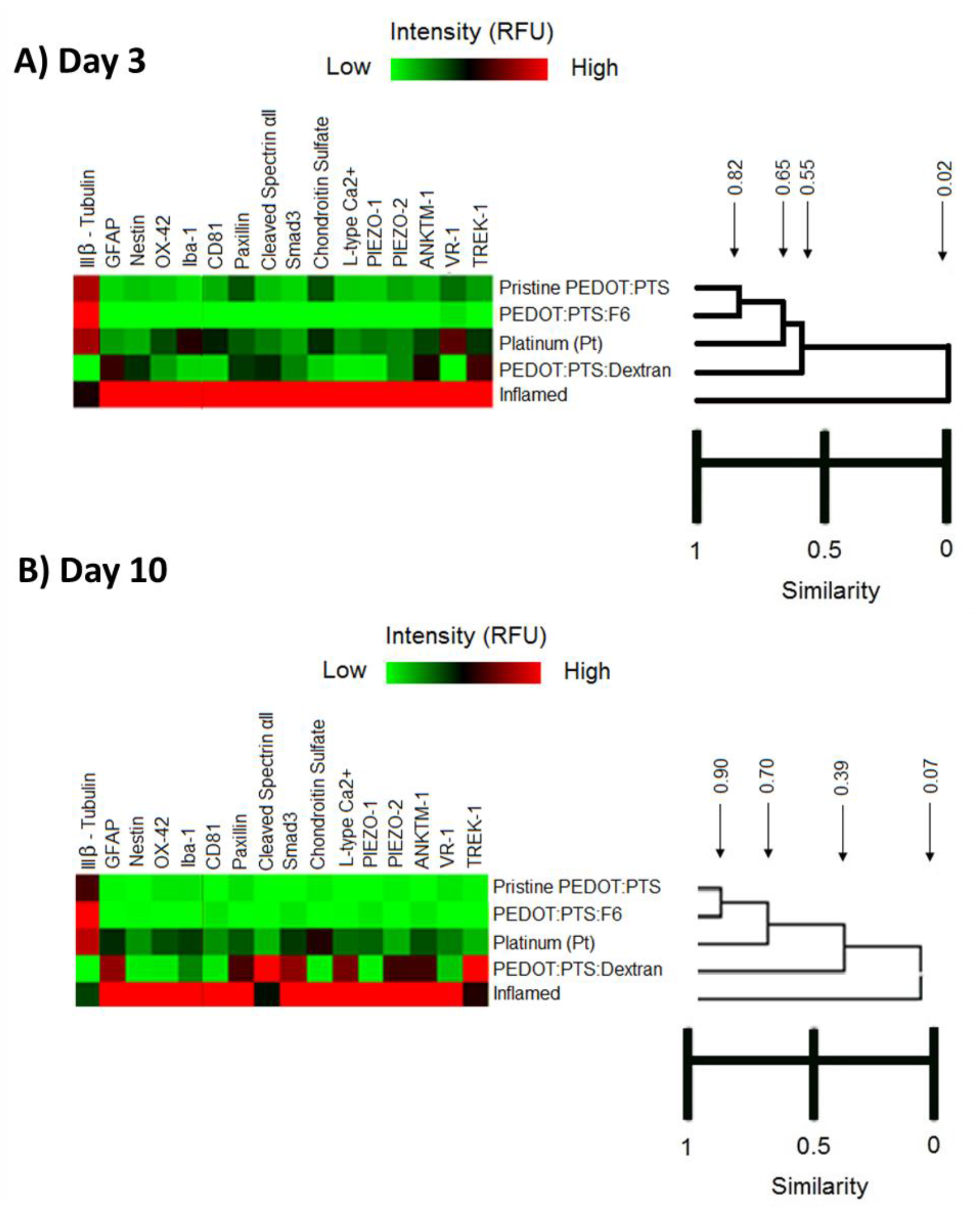
Unsupervised clustering of all experimental and control groups and inflamed control, respectively. This analysis clearly defined interestingly hierarchical clusters between experimental sulphated groups: the chemical sulphated pristine PEDOT:PTS coated electrodes and the biological modified PEDOT:PTS:F6 coated electrodes with the heparan sulphate mimetic-F6, and between controls groups with non-sulphated patterns: control material bare platinum, the negative control for sulphation PEDOT:PTS:Dextran coated electrodes and tissue culture plastic as inflamed control, at both days three A) and ten B) respectively. PEDOT:PTS:F6 coated electrodes showed a remarkable intensity downregulation of reactive astrocytes, microglia and calcium influx markers with an important upregulation of neural response compared to all groups and controls at day three and day ten, respectively. N = 3, 18 data points.

This analysis defined groups with hierarchical similarities between cells cultured on experimental sulphated groups, the chemically sulphated pristine PEDOT:PTS coated electrodes and the biological modified PEDOT:PTS:F6 coated electrodes with F6, followed by controls groups with non-sulphated patterns, i.e. control Pt, the negative control for sulphation PEDOT:PTS:Dextran coated electrodes and finally tissue culture plastic as inflamed control, at both three and ten days.

By day three, PEDOT:PTS:F6 coated electrodes induced a marked downregulation in the expression of pro-gliosis proteins relative to all experimental, and inflamed controls. Specifically reductions were observed in the expression of OX42/Iba1 (microglia) [166], GFAP and Nestin (reactive astrocytes) [167, 168], gliosis specific markers such as cleaved spectrin αII [169],Smad3 [170], Chondroitin Sulphate [171] and CD 81 [172, 173], as well as from the changes of transient calcium seen by the expression of L-type Ca^2+^ [174], PIEZO-2 [175], K^+^ channels such as TREK 1 [176], and in the low intensity of proteins linked to inflammation such as VR1 [177] and ANKTM1 [178]. Interestingly, by day three a high expression of the neural marker III β Tubulin was observed relative to all experimental, controls and inflamed groups (***Figure 17*A**). By day ten although PEDOT:PTS:F6 coated electrodes shared a 90% protein expression similarity with the pristine PEDOT:PTS coated electrodes, a significant increase in the expression of β-tubulin was observed, indicating a relative increase in the presence of VM neurons on PEDOT:PTS:F6 functionalised electrodes (***Figure 17*B**). These findings are consistent with the noticeable neural survival observed on the PEDOT:PTS:F6 coated electrodes (***Figure 14*B**) and with the suggested neurotropic effect of F6 relative to all experimental, control and inflamed groups. Further and owing to the multifactorial markers of the hallmark of gliosis evaluated here with the microarray, the results also allude to the overall low inflammation profile offered by the PEDOT:PTS:F6 functionalised electrodes F6.

In contrast, the negative control for glycan sulphation, the PEDOT:PTS:Dextran coated electrodes, exhibited the lowest intensity of the neural marker, III β Tubulin compared to all experimental, and control groups except the inflamed control, at day three and further maintained by day ten. Interestingly, PEDOT:PTS:Dextran coated electrodes induced a GFAP expression profile similar to that of VM cells cultured under inflamed control conditions, but induced a marked differential intensity in the expression of the K^+^ ion channel TREK 1 relative to all experimental and control groups. This suggest a disruption of astrocyte K^+^ homeostasis, which has been shown to help set the negative resting membrane potential of astrocytes and regulate astrocyte reactivity [179], an effect that is reflected in the over-proliferation of astrocytes observed on these coated electrodes relative to PEDOT:PTS:F6, pristine PEDOT:PTS coated electrodes and Pt controls (***Figure 14*B**).

Coupled with this, marked upregulations in the expression of cleaved spectrin αII relative to the inflamed control group [169], together with a high expression in Smad3 [170], and paxillian, an adhesion marker used for the healthy stellation of astrocytes [180–182], indicated an overall significant influence on reactive astrocyte adhesion on PEDOT:PTS:Dextran electrodes relative to PEDOT:PTS:F6 and pristine PEDOT:PTS coated electrodes and Pt control electrodes. The further reduced expression of OX42/Iba1 and VR1, markers for microglia [183], and increased expression of ion channel proteins (L-type V [174] and PIEZO-2 [152]) by day ten, may reflect the overall pro-inflammatory nature of PEDOT:PTS:Dextran coated electrodes relative to PEDOT:PTS:F6, pristine PEDOT:PTS coated electrodes and Pt control.

Experimentally, it has been demonstrated that the HS mimetic F6, when incorporated in a PEDOT:PTS polymer matrix, is able to enhance neuron survival and support neural outgrowth while maintaining a low inflammatory and low gliosis micro-environment *in vitro*.

In order to assess the effects of F6 on the glycosylation profile of labeled protein preparations VM populations, a lectin binding assay employing 26 lectins was conducted following three and ten days of culture. Relative fluorescence intensity (RFU) values for lectins that exceeded approximately five times the average median background (1,000 RFU) were considered to have undergone glycan binding and were contributed to the glycosylation pattern.

A marked binding intensity with the lectins SNA-I and AAL was observed in all experimental and control groups (***Figure 18***), which indicated the presence of α-(2,6)-linked sialic acid and fucosylation, respectively (***Table 2***). AAL has a binding specificity with high affinity for α −(1,6)-linked fucose (Fuc) residues which are found on the chitobiose core of N-linked oligosaccharides, and can also bind with much lower affinity to α −(1,3)- and α −(1,2)-linked Fuc [184, 185] (***Table 2***). Interestingly, α −(2,3)-linked sialic acid has been previously shown to be the most abundant sialic acid linkage present in total brain tissue protein [186]. The predominance of α-(2,6)-linked sialic acid in the VM cell protein preparations may be characteristic of the particular type compared to total tissue.

**Figure 18.**
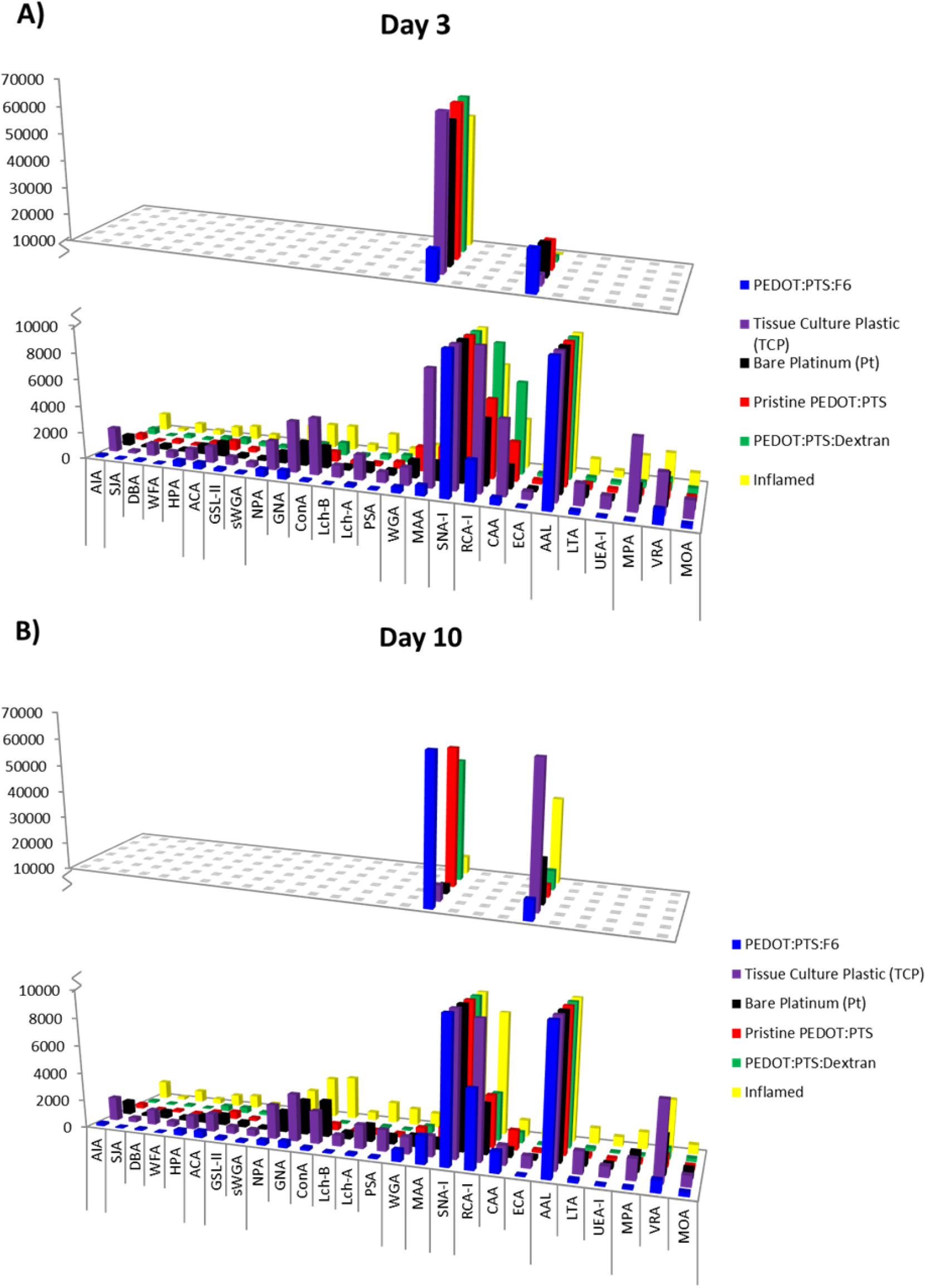
Histograms representing the differences in recognition of printed lectins by each of the experimental groups and controls at day three A) and day seven B) respectively. Microarray experiments were carried out using three replicate slides. The subarrays were printed with each lectin following the order detailed in Figure 4.2 and incubated with fluorescently labelled protein preparations. Results are average ± standard error. N = 3, 18 data points

At day three VM cells cultured on all experimental and control groups exhibited glycosylation profiles with a high binding affinity to SNA-I and AAL, which presented differential low binding affinity on inflamed and tissue culture plastic controls and on PEDOT:PTS:Dextran coated electrodes relative to VM cells culture on Pt controls and pristine PEDOT:PTS, with the highest binding observed in VM cells cultured on PEDOT:PTS:F6 coated electrodes (***Figure 18*A**). Further, VM cells cultured on PEDOT:PTS:F6 functionalised electrodes, exhibited almost equally intense binding for SNA-I and AAL, with SNA-I binding intensities significantly lower than all other experimental and control groups. Critically, glycoprotein sialylation, is known to play a role in modulating the excitability of voltage-gated ion channels, and in mediating cellular adhesion [187, 188]. Moreover, sialic acids can function as ligands for sialic acid-specific receptors, such as siglecs and selectins, facilitating cellular communication and signaling with roles in inflammation [187, 189] and cell fate [190].

Fucosylation, as identified through high binding affinity to AAL has a roles in mediating interactions with selectins and α-(1,6)-linked fucosylation is a common modification of the rat brain N-linked oligosaccharides [186]. This motif has been shown to play an important role in regulating cell neurite formation [191] while a loss of α-(1,6)-fucosylation has been shown to decrease hippocampal potentiation [192]. Interestingly, the observed higher binding affinity for α-(1,6)-fucosylation (AAL) in VM cells cultured on PEDOT:PTS:F6 coated electrode relative to all experimental and control groups may suggest on the early trophic effect of F6.

By day ten all experimental conditions were associated with a reduction in SNA-I binding efficiency relative to day three, except for VM cells cultured on PEDOT:PTS:F6 coated electrodes, which exhibited increased SNA-I binding (***Figure 18*B**). This binding intensity was comparable to all other experimental and control groups at day three. This result may suggest on the role of glycoprotein sialylation in modulating the excitability of voltage-gated ion channels [187, 188]. The promotion of sialic acid expression through electrode functionalisation with the synthetic HS mimetic may be as a response of neuronal control and network excitability through electrical signalling, providing adequate conditions for the appropriate function of neurons and synapses [193, 194]. This data can be further correlated with the downregulation of voltage-gated ion channels observed by the VM cells cultured on the functionalised electrodes, followed by pristine PEDOT:PTS coated electrodes compared to all controls at day ten (***Figure 17*B**). In turn, impairment of K^+^ buffering currents by the altered high expression of TREK-1 channels by astrocytes on PEDOT:PTS:Dextran coated electrodes by day ten, may contribute, particularly to the high sialic acid content observed on these electrodes [195–197]. Results that suggest on the importance of orchestrated variable sets of ion channels for homeostasis [198]. Furthermore, the maintained expression of AAL binding for 1,6-fucosylation may suggest to an overall temporal-regulation of intracellular signalling and neural homeostasis offered by the presence of the HS mimetic.

### 2.3 Conclusion

In the search for biomimicry of the neuroelectrode interface, and with an ultimate goal of mitigating electrode deterioration via reactive host cell response and glial scar formation, PEDOT:PTS neural coating were functionalised with a heparan mimetic as a biological dopant for the first time.

The use of F6 as a dopant within PEDOT:PTS electroactive coatings significantly reduced the electrochemical resistance and increased the charge storage capacity of Pt/Ir electrodes, while maintaining electrode stability. Together, neuroprotective and neurotrophic effects were observed *in vitro* with a significant reduction in the inflammatory and gliosis profile noted in a complex primary mixed cell VM culture accompanied by neurogenesis and neurotogenesis of the mesencephalic neurons. These trophic characteristics can be explained by the potentiation of synthetic HS mimetic-F6, in the binding of FGF-2 and VEGF growth factors to the functionalised peri-electrode region.

Bio-functionalisation of neural electrodes with PEDOT:PTS:F6 coatings show promise as a functional approach to promote the integration of implanted electrodes through the attenuation of glial scar response and through promoting a neuroprotective response *in vitro*. This work further promotes the exploration of further glycan mimetics to potentiate matrix-therapeutics strategies at the neural-tissue interface.

